# Increased Replication of Dissimilatory Nitrate-Reducing Bacteria Leads to Decreased Anammox Bioreactor Performance

**DOI:** 10.1101/534925

**Authors:** Ray Keren, Jennifer E. Lawrence, Weiqin Zhuang, David Jenkins, Jillian F. Banfield, Lisa Alvarez-Cohen, Lijie Zhou, Ke Yu

**Author notes:** These authors contributed equally to the work (Ray Keren, Jennifer E. Lawrence). Corresponding authors Lijie Zhou; Ke Yu. Email of Authors: Ray Keren; Jennifer E. Lawrence; Weiqin Zhuang; David Jenkins; Jillian F. Banfield; Lisa Alvarez-Cohen.

## Abstract

**Background:** Anaerobic ammonium oxidation (anammox) is a biological process employed to remove reactive nitrogen from wastewater. While a substantial body of literature describes the performance of anammox bioreactors under various operational conditions and perturbations, few studies have resolved the metabolic roles of their core microbial community members.

**Results:** Here, we used metagenomics to study the microbial community of a laboratory-scale anammox bioreactor from inoculation, through a performance destabilization event, to robust steady-state performance. Metabolic analyses revealed that nutrient acquisition from the environment is selected for in the anammox community. Dissimilatory nitrate reduction to ammonium (DNRA) was the primary nitrogen removal pathway that competed with anammox. Increased replication of bacteria capable of DNRA led to the out-competition of annamox bacteria, and the loss of the bioreactor’s nitrogen removal capacity. These bacteria were highly associated with the anammox bacterium and considered part of the core microbial community.

**Conclusions:** Our findings highlight the importance of metabolic interdependencies related to nitrogen- and carbon-cycling within anammox bioreactors and the potentially detrimental effects of bacteria that are otherwise considered core microbial community members.

## Background

Anaerobic ammonium oxidizing (anammox) bacteria obtain energy from the conversion of ammonium and nitrite to molecular nitrogen gas (N_2_) [1]. Currently, the only bacteria known to catalyze this process are members of the phylum *Planctomycetes* [2, 3], none of which have been isolated [3, 4]. In practice, anammox bacteria are employed in an eponymous process in combination with the partial nitritation (PN) process to remove ammonium from nitrogen-rich wastewaters. First, in PN, approximately half of the ammonium in solution is aerobically oxidized to nitrite. Second, in anammox, both ammonium and nitrite are anaerobically converted to N_2_ [5, 6]. The PN/anammox (i.e., deammonification) process is beneficial because it consumes 60% less energy, produces 90% less biomass, and emits a significantly smaller volume of greenhouse gas than conventional nitrogen removal by nitrification and denitrification processes [7]. To date, over 100 full-scale deammonification process bioreactors have been installed at municipal and industrial wastewater treatment plants across the globe [8].

Within an engineered environment, anammox bacteria have very low growth rates and can easily become inhibited by fluctuating substrate and metabolite concentrations [9, 10]. When these two limitations are coupled, recovery from an inhibition event can take up to six months (which is unacceptably long for municipalities that must meet strict nitrogen discharge limits) [11]. Furthermore, these problems are compounded by a cursory understanding of the microbial communities that exist alongside anammox bacteria. A deeper understanding of the complex interactions occurring among bacterial species in an anammox bioreactor is required for the broad application of the deammonification process for wastewater treatment.

Previous research has suggested that a core microbial community exists within anammox bioreactors [12–16]. In the majority of studied bioreactors, uncultured members of the phyla *Bacteroidetes*, *Chloroflexi*, *Ignavibacteria*, and *Proteobacteria* have been identified alongside *Planctomycetes*, the phylum that contains anammox bacteria. These phyla have primarily been identified through 16S rRNA gene studies, so their interplay with anammox performance has not yet been fully elucidated [12–16]. From their taxonomic identity and performance studies, it is assumed that the additional phyla compete for nitrite and cooperate to transform (i.e. dissimilatory nitrate reduction to ammonium - DNRA) and remove (i.e. denitrifiers) nitrate, a product of anammox metabolism [17–19].

Here, we illuminate deeper metabolic relationships between an anammox bacterium, *Brocadia*, and its supporting community members during the start-up and operation of a laboratory-scale anammox bioreactor. We begin by analyzing the formation of the “anammox community” through a combination of genome-centric metagenomics and 16S rRNA gene sequencing. Metabolic features of positively enriched bacteria are compared to negatively enriched bacteria during the start-up process. Next, we focus our investigation on an anammox performance destabilization event that was driven by microbial interactions. Last, we conduct a comparative analysis of our anammox community to similarly-studied anammox communities [18, 20] to highlight the broader relevance of our results. To our knowledge, this is the first time-series-based study to link anammox metagenomic insights and community composition to anammox bioreactor functionality [21]. Our findings bolster the fundamental, community-level understanding of the anammox process. Ultimately, these results will enable more comprehensive control of this promising technology and facilitate its widespread adoption at wastewater treatment plants.

## Results

### Bioreactor performance

The performance of a laboratory-scale anammox anaerobic membrane bioreactor (MBR) (described in methods) was tracked for 440 days from initial inoculation, through several performance crashes, to stable and robust anammox activity (Figure 1). Performance was quantified by nitrogen removal rate (NRR, g-N L^-1^ d^-1^) and effluent quality (g-N L^-1^ d^-1^). Bioreactor performance generally improved over the first 103 days of operation. At this point, the hydraulic residence time (HRT) was reduced from 48 to 12 hours and influent nitrogen concentrations were reduced to maintain a stable loading rate. Additional seed biomass from a nearby pilot-scale deammonification process was added on Day 145 following a performance crash and bioreactor performance improved, enabling influent ammonium and nitrite concentrations to be steadily increased until the NRR approached 2 g-N L^-1^ d^-1^. On Day 189 the bioreactor experienced a technical malfunction and subsequent performance crash, identified by a rapid decrease in the NRR and the effluent quality. On Day 203, the bioreactor was again amended with a concentrated stock of seed biomass and the NRR and the effluent quality quickly recovered. Influent ammonium and nitrite concentrations were again increased until the NRR reached 2 g-N L^-1^ d^-1^.

**Figure 1.**
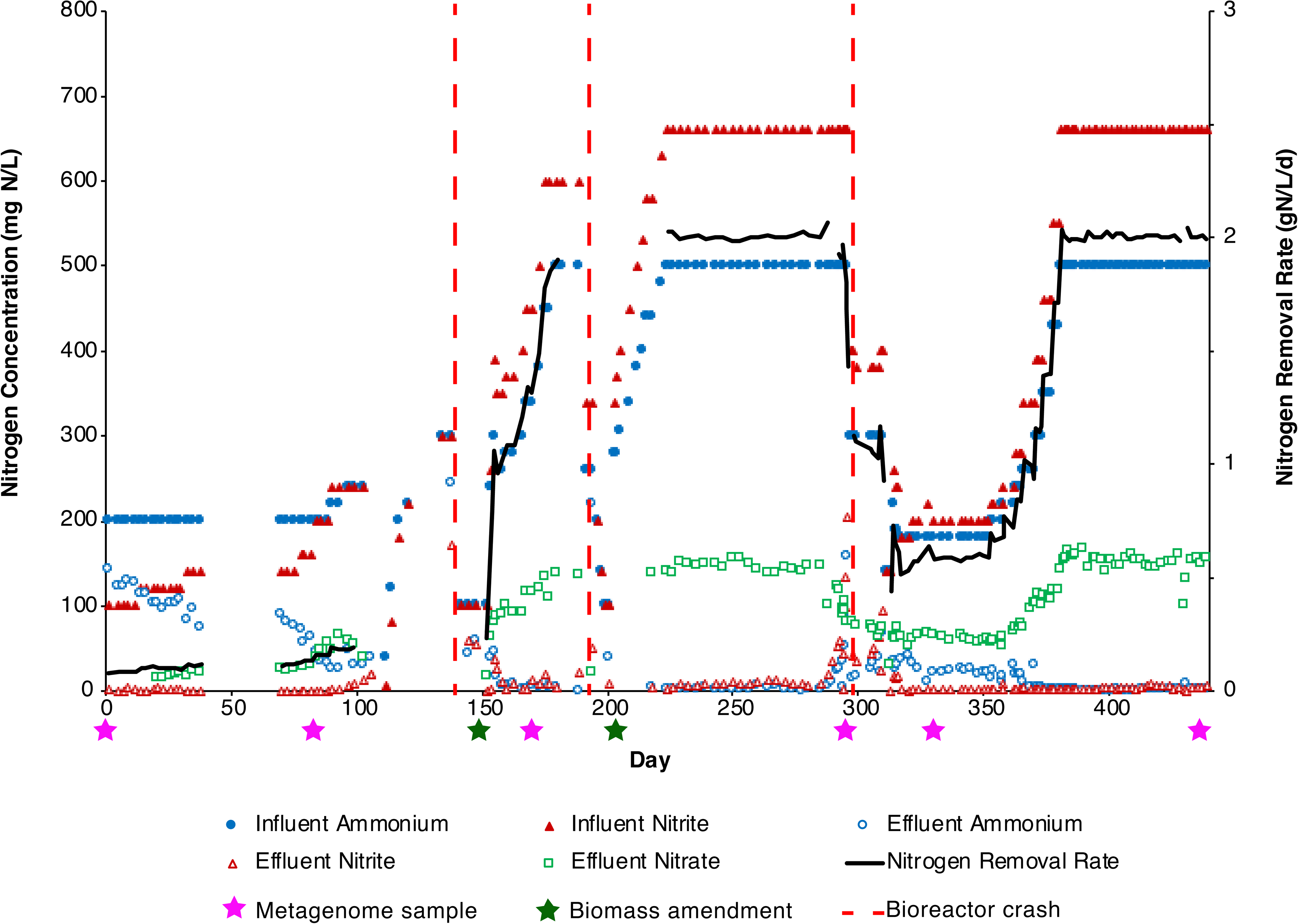
Performance of the anaerobic membrane bioreactor. Influent and effluent concentrations of ammonium, nitrite, and nitrate (all as N) (primary y-axis) within the anaerobic membrane bioreactor performing anammox monitored over a period of 440 days. The influent did not contain nitrate, so influent nitrate is not plotted. The nitrogen removal rate (NRR), is plotted against the secondary y-axis. Sampling time points for metagenomes are indicated with purple stars below the x-axis. Biomass amendments are indicated with green stars below the x-axis. Bioreactor crashes (either mechanical or biologically driven) are marked with a dashed red line.

The bioreactor subsequently maintained steady performance for approximately 75 days, until Day 288, when effluent concentrations of ammonium and nitrite unexpectedly began to increase and nitrate concentrations disproportionately decreased. Seven days later, the NRR rapidly plummeted. No technical malfunctions had occurred, indicating that the destabilization of the anammox process may have been caused by interactions among its microbial community members. At the time, the cause of the performance decline was not understood, so the bioreactor was not re-seeded with biomass. After 50 days of limited performance, concentrations of copper, iron, molybdenum, and zinc in the bioreactor influent were increased based on literature recommendations [22–25] and the NRR rapidly recovered. Stable and robust bioreactor performance was subsequently maintained.

### Metagenomic sequencing and binning

Whole community DNA was extracted and sequenced at six time points throughout the study: Day 0 (D0), for inoculant composition; Day 82 (D82), during nascent, positive anammox activity; Day 166 (D166), three weeks after an additional biomass amendment; Day 284 (D284), after a long period of stable and robust anammox activity and just before the bioreactor performance was destabilized; Day 328 (D328), in the midst of the performance destabilization period; and Day 437 (D437), during mature, stable, and robust anammox activity.

From all samples, 337 genomes were binned, 244 of which were estimated to be >70% complete by checkM [26]. The genomes were further dereplicated across the six time points into clusters at 95% average nucleotide identity (ANI). This resulted in 127 representative and unique genomes (Table S1) that were used for all downstream analyses. Mapping showed an average read recruitment of 76% to the representative genomes (Table 1). The number of genomes present at each time point (using threshold values of coverage > 1 and breadth > 0.5) ranged from 60 (D437) to 103 (D166). In addition, nine strains were detected that differed from the representative genome by 2% ANI (Additional file 1, Table S2). Except for the anammox bacterium, which is referenced at the genus level (*Brocadia*), all representative genomes are referenced at the phylum level.

**Table 1.** Read counts to representative genomes across time points.

| Time point (days) | Total read number | Mapped read number | % of reads mapped | Number of genomes detected* |
| --- | --- | --- | --- | --- |
| 0 | 55398280 | 40291503 | 72.73 | 92 |
| 82 | 62544544 | 43877427 | 70.15 | 82 |
| 166 | 60931806 | 46030350 | 75.54 | 103 |
| 284 | 56282006 | 48644523 | 86.43 | 68 |
| 328 | 127048582 | 95132145 | 74.88 | 87 |
| 437 | 119945232 | 93737087 | 78.15 | 60 |

\* number of representative genomes based on threshold of >1 coverage and > 0.5 breadth

### Community structure and temporal dynamics

As both internal and external factors can work in combination to affect the structure of a bioreactor community, we hypothesized that different groups of bacteria (i.e., sub-communities) would be associated with different phases of the bioreactor’s lifespan. To test for grouping, all genomes were pairwise-correlated (Figure 2A). The resulting heatmap revealed four distinct clusters (Groups A-D). Group A was the largest, with 52 genomes, while Groups B-D had 25, 24, and 26 genomes respectively (Table S3).

**Figure 2.**
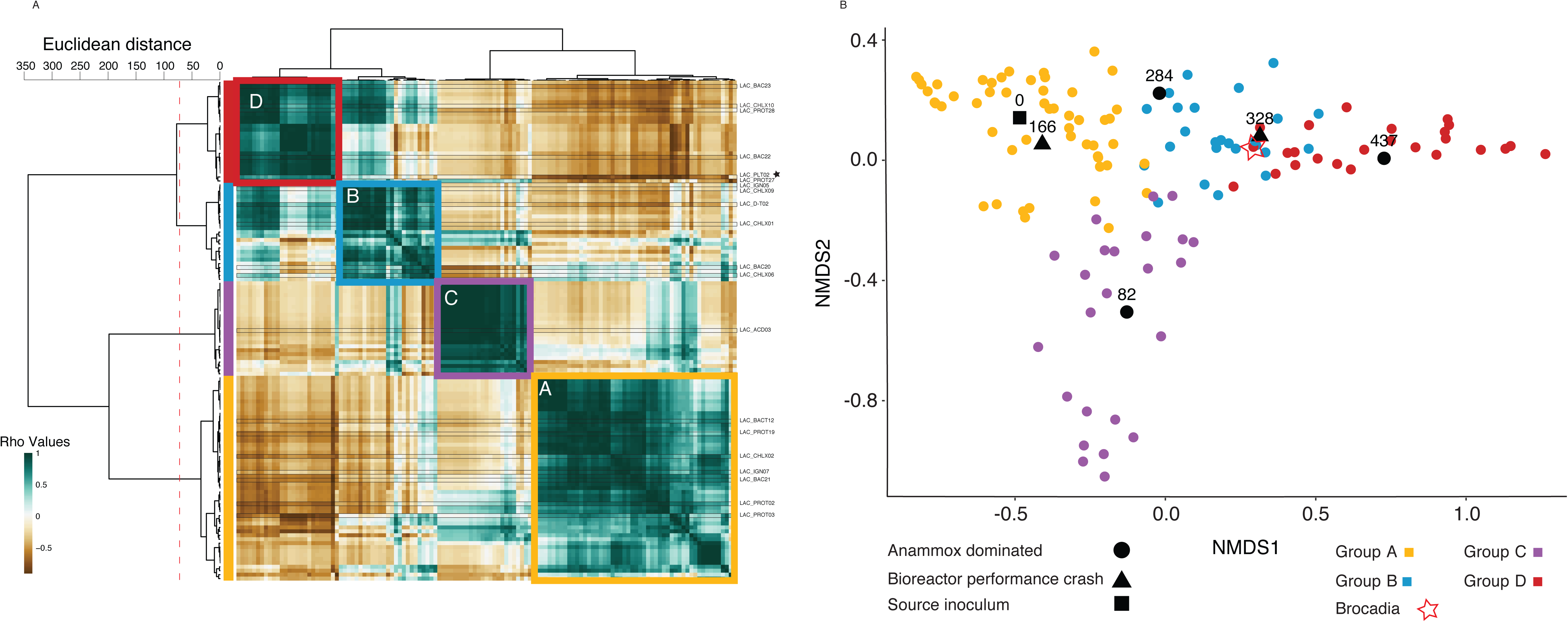
Analysis of the bioreactor community clustering by the estimated relative abundance of the bacteria. (A) clustering heatmap of bacteria based on pairwise cross correlations for the six time points (matrix values are Rho values). Color scales mark high positive correlation in green and negative correlation in brown. The row and column dendrograms are identical (consisting of the genomes). The row dendrogram shows the calculated distance between the clusters with a dashed red line marking the split to clusters. Colored squares as well as bars to the left of the heat map show relative abundance based grouping: Yellow - Group A, Blue - Group B, Purple - Group C, Red - Group D. A black star to the right of the heatmap marks the anammox bacterium (*Brocadia*). (B) Two-dimensional nMDS projection of bacteria and time points, showing the association of the bacteria (and relative abundance groups to certain time points). Each colored dot represents the centroid of a bacterium, with colors matching the relative abundance group. Black marks represent the centroid of the time points, while the shape represents the state of the bioreactor: Circle - anammox dominated, Triangle - bioreactor crash (either mechanical or biological), Square - time zero. The location of *Brocadia* is marked with a red star.

To better examine the clustering of genomes in relation to the bioreactor’s lifespan, we ran nonmetric multidimensional scaling (nMDS) analyses on the genomes’ relative abundance data (Figure 2B). The nMDS projection revealed that genome groups were strongly associated with specific time points: Group A was associated with the inoculant biomass at D0 and D166, while Group C was associated with the nascent anammox community at D82. Group B was associated with the times of destabilized anammox performance (Days 284-328), and Group D was associated with the mature, stable anammox community at D437. *Brocadia* is a part of Group D, although its location on the nMDS projection is skewed to the left because of its high relative abundance throughout the majority of the bioreactor’s lifespan. Because the nascent anammox community was amended with additional biomass, we could not resolve a linear trajectory for the microbial community between the initial and final states. Nevertheless, Groups B and D shared many similarities, and the majority of the genomes associated with Group B were still present in the bioreactor on D437.

To further resolve the relative abundances of Groups A-D over the bioreactor’s lifespan, 16S rRNA genes from genomes were combined with direct 16S rRNA sequencing data, then clustered into operational taxonomic units (OTUs). Of the 127 representative genomes, 34 contained a 16S rRNA sequence that was successfully clustered to an OTU. The matching OTUs represented 55% of the total 16S rRNA reads at day 9, but quickly increased to an average representation of 86%. The OTUs that match assembled genomes were grouped (Figure 2) and their relative abundance summed (Figure 3A). The matching genomes comprised 18/52, 10/25, 3/24, and 7/26 of Groups A-D respectively. The matching OTUs represented 55% of the total 16S rRNA reads at day 9, but quickly increased to an average representation of 86%.

**Figure 3.**
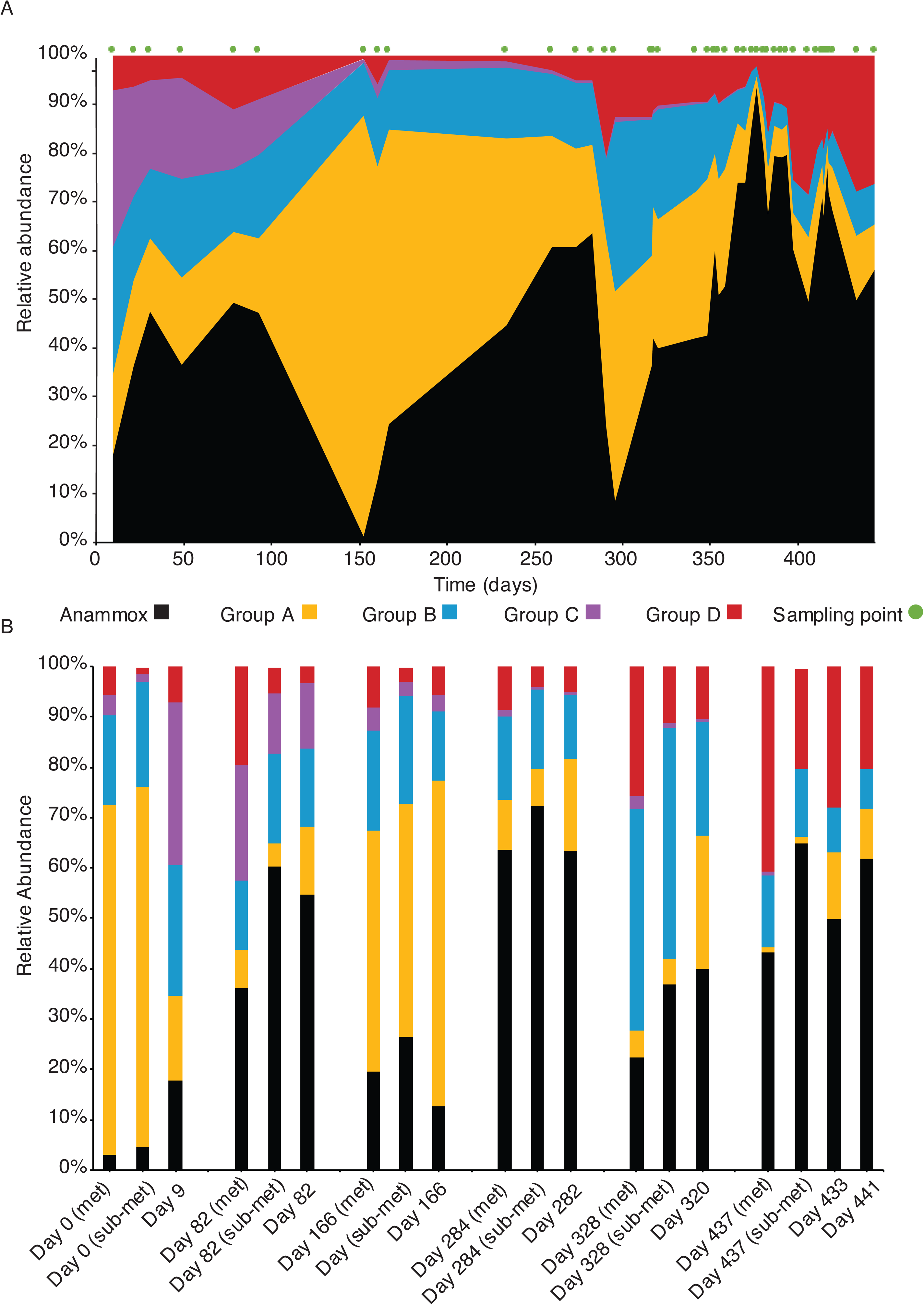
Relative abundances of bacterial groups over the lifespan of the bioreactor. **(A)** Relative abundances of bacterial groups A-D, based on 16S rRNA OTUs that have a match to a draft genome. Group colors are matched by the analysis in Figure 2, with the exception of *Brocadia*, which was removed from group D and is depicted in black. Green points above the graph show timepoints where the community was sampled. (B) Comparison of relative abundances of the different groups at the same time (or the closest time). Relative abundance was calculated for groups based on all the draft genomes (marked met on the x-axis), the subset of genomes with matched to the 16S rRNA OTUs (marked sub-met on the x-axis) or the 16S rRNA OTUs (only day is marked on x-axis).

Group A was dominant on Day 0, but rapidly decreased in abundance by the first 16S rRNA gene sequencing time point on Day 9 (Figure 3B). Group A was again dominant after the addition of new inoculum on Day 145 (Figure 3A). Group B (and to a certain extent Group A) became dominant just before Day 300 when the anammox performance was destabilized.

To check the accuracy of the 16S rRNA matching to the metagenomic data, we compared the relative abundances of Groups A-D across three data sub-sets (Figure 3B): all metagenomes (marked ‘met’ on the x-axis), only metagenomes with matching 16S rRNA OTUs (marked ‘sub-met’ on the x-axis), and the 16S rRNA OTUs. Overall, the three datasets were compatible, with slight variations in over/under-estimations of particular Groups. In comparison to the metagenomically-derived data, the 16S rRNA data tended to overestimate the relative abundance of Group A and underestimate the relative abundance of Group D. A large fraction of the *Chloroflexi* in Group D were not matched to 16S rRNA OTUs, so the underestimation was consistent with expectations.

For all subsequent analyses, we split the representative genomes into two groups (Table S3): those that are associated with the mature anammox community at D437 (Anammox Associated, AA), and those that are not (Source Associated, SA). The AA community includes all of the genomes that are present at D437, while the SA community includes the rest of the genomes that are not present at D437. Some of these genomes are associated with the sludge amendments, and some are associated with the nascent anammox community; at no point is there a community exclusively comprised of SA genomes.

### Metabolic profiles

For the purpose of analyzing the metabolic potential of the microbial community, we evaluated only genomes with > 70% completeness (n = 88) [26]. Using Hidden Markov Model (HMM) searches of the Kyoto Encyclopedia of Genes and Genomes (KEGG) database, we checked for the presence of genes based on their KEGG Orthology (KO) number, and calculated KEGG module completeness [28, 29]. The genomes were clustered by KO presence/absence (Figure S1) and their module completeness (Figure 4). Clustering by the two methods resulted in similar groupings.

**Figure 4.**
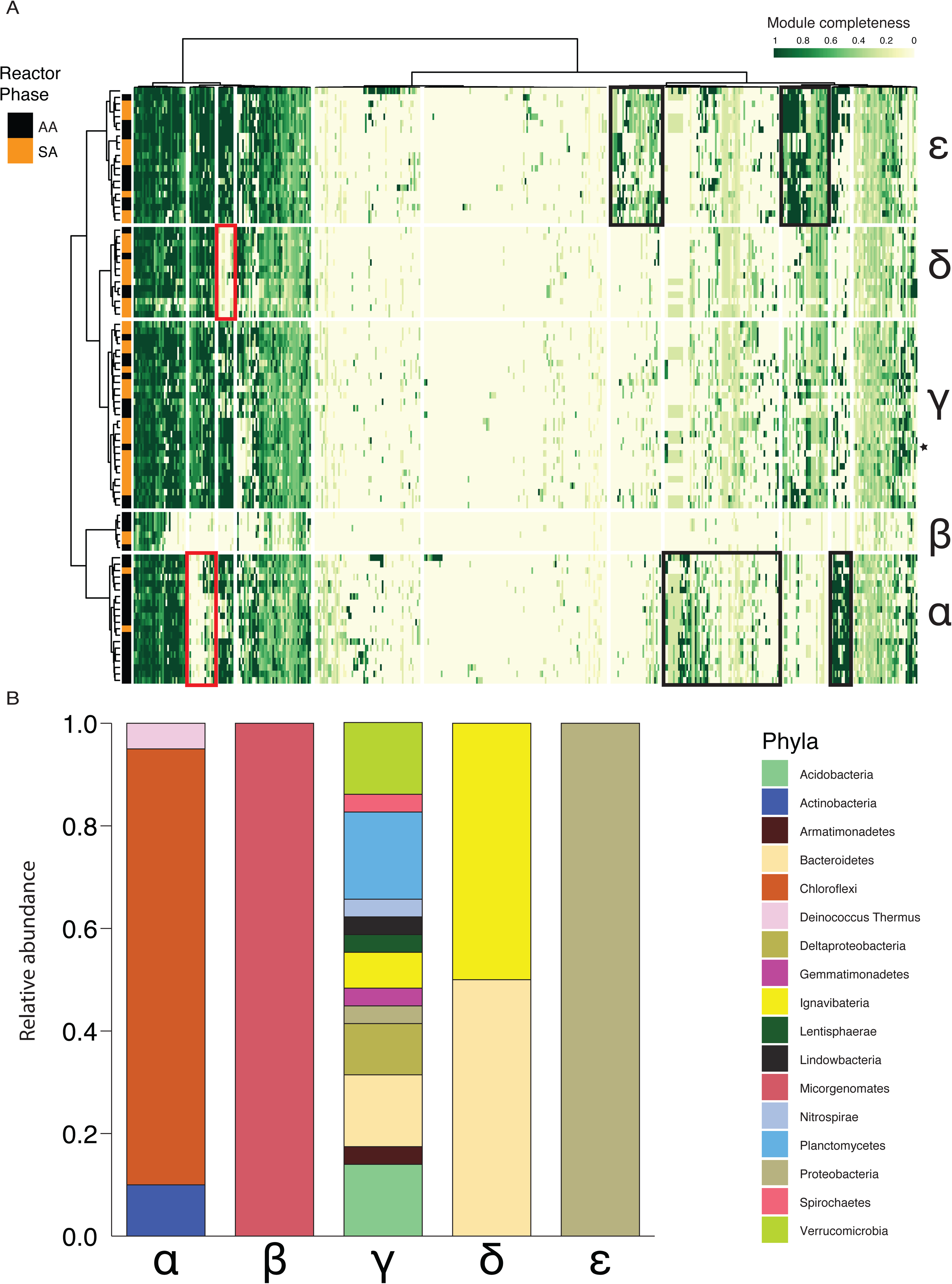
Metabolic profiling of bacterial community based on KEGG module completeness. (A) heatmap showing clustering of genomes (rows) by their KEGG module completeness (columns). Completeness ranges from 1 (green) to zero (white). The heatmap is based on a Euclidean distance matrix and clustering with the ward.D method. Genome clustering resulted in 5 clusters (groups α-ε). Rectangles on the heatmap mark module blocks that differentiate the genome groups. Black rectangles on heatmap show module blocks that have increased completeness in a group of bacteria (compared to the others), and red rectangles show decreased completeness. Marks to the left of the heatmap show the division of AA and SA bacteria. A black star to the right of the heatmap marks *Brocadia*. B) Relative abundance by phyla of members in each metabolic cluster.

Module clustering resolved five groups (□, β, □, δ, ε) (Figure 4A and Table S3). Groups □ and β contained more anammox-associated genomes (90% and 60%, respectively), while groups □, δ, and ε contained 65%, 70% and 60% of source-associated genomes, respectively. The clustering was also strongly influenced by bacterial taxonomy (Figure 4B). Group (□ was composed solely of Gram (+) bacteria, primarily *Chloroflexi*. Group β was composed of Candidate Phyla Radiation (CPR) bacteria, *Microgenomates*. This group of bacteria has reduced genomes and metabolism [30], and thus has an unknown effect on the community metabolism. Group □ was composed entirely of Gram (-) bacteria from a wide range of phyla, and includes the anammox bacterium *Brocadia*. Group δ was composed of *Ignavibacteria* and *Bacteroidetes*, but only the *Ignavibacteria* from Group δ were associated with the AA group. Accordingly, further analysis of Group δ refers only to the *Ignavibacteria*. Group ε was composed entirely of *Proteobacteria*.

Based on the KEGG module clustering, we reconstructed the representative metabolisms of the five groups (Figure 5). We used a module completeness threshold of 67% per genome and considered it representative if it was complete in >50% of the group’s members. Group δ was not represented since it diverged from group □ by auxotrophies in several modules (Figure 4A, red rectangle). The *Brocadia* metabolism is shown in Figure S2.

**Figure 5.**
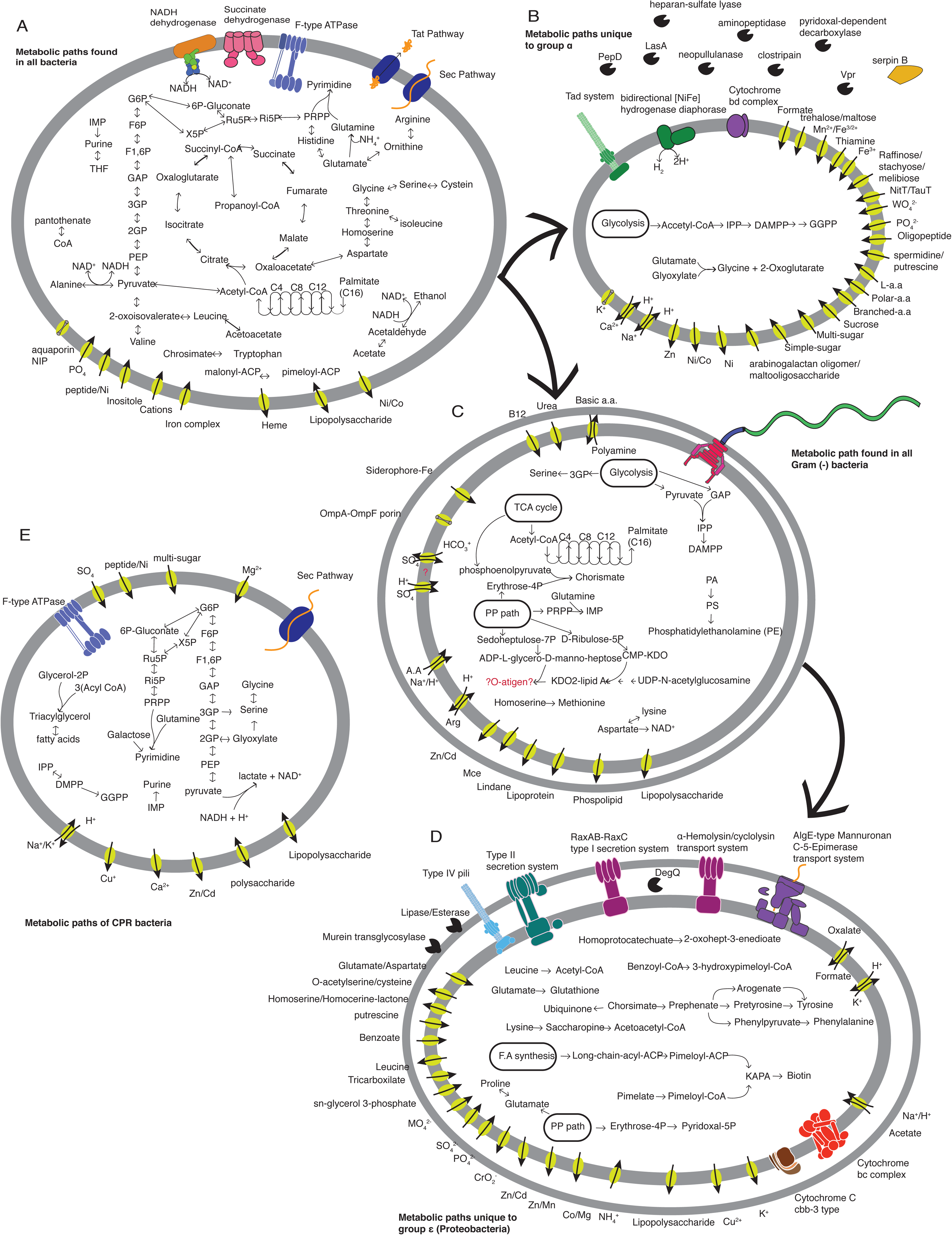
Representative metabolic maps of bacterial groups in the bioreactor. To prevent redundancy, the metabolism is presented in a nested approach with each panel showing only paths unique to the relevant metabolic group. Two exceptions are group β (all detected paths are shown), and group δ. The latter is not presented here since it shares all paths with group □ and only differs by auxotrophies. (A) Metabolic map of paths that are common to all bacteria in the bioreactor (except *Microgenomates* and *Brocadia* sp.). The vast majority of bacteria in the bioreactor are heterotrophs, capable of carbohydrate-based metabolism (glycolysis, pentose phosphate pathway) and amino acid-based metabolism. Some bacteria can respire oxygen, and can also ferment (acetate/alanine). (B) Paths unique to group □. These bacteria have genes for hydrogen oxidation, supporting anaerobic growth, as well as genes for oxidative phosphorylation with cytochrome BD complex. These bacteria have a cassette of extracellular proteases and decarboxylases, paired with a wide array of transporters. Theyare also potentially capable of synthesizing long chain isoprenoids. (C) Paths found in Gram (-) bacteria (groups □, δ, and ε). Most paths are related to fatty acid and lipid synthesis. Several important precursors (chorismate and IMP) can potentially be synthesized by these bacteria. Motility is also a common feature in these bacteria (via a flagellar motor) (D) Unique paths of group ε (*Proteobacteria*). This group has the potential to synthesize multiple vitamins and cofactors (biotin, pyridoxal, glutathione, etc.), as well as several amino acids (tyrosine, phenylalanine, proline). Another unique feature is the multiple secretion systems present in the bacteria. (E) Metabolic profile of CPR bacteria (*Microgenomates*). These bacteria are obligate anaerobes that ferment pyruvate. They can only utilize carbohydrates as their carbon source. Some of the bacteria in this group might also be able to synthesize long chain isoprenoids, in the same path as group □.

While module completeness was used for most of the analyses, in several cases it was not sufficient (e.g., overlap between modules, no module for path). For oxidative phosphorylation, fermentation, carbon fixation, several amino acid synthesis pathways, and nitrogen metabolism, we analyzed gene presence manually. For anammox, four additional HMMs were added: hydrazine synthase subunit A (*hzsA*), hydrazine oxidoreductase subunit A (*hzoA*), and nitrite oxidoreductase subunits *nrxA* and *nrxB* [31]. For the latter, the similarity of the gene to the nitrate reductase *narGH* was taken into consideration.

With the exception of two CPR bacteria, all of the genomes in the bioreactor contained genes encoding assimilation of ammonia into glutamate (Figure 6). More than half (49) of the bacteria could potentially reduce nitrate, and the same number had the genes needed to further reduce nitrite to nitrogen monoxide (NO); however, only 26 bacteria had the genes to do both steps. The remaining steps of denitrification were encoded in an even smaller number of genomes. The *nrxAB* gene was only identified in two genomes, one of which was *Brocadia.* One-step DNRA was identified in 22 genomes. While the number of genes encoding ammonia assimilation and nitrate reduction to nitrite were fairly similar in the AA and SA groups’ genomes, DNRA was more common in the AA genomes and denitrification beyond nitrite in the SA genomes.

**Figure 6.**
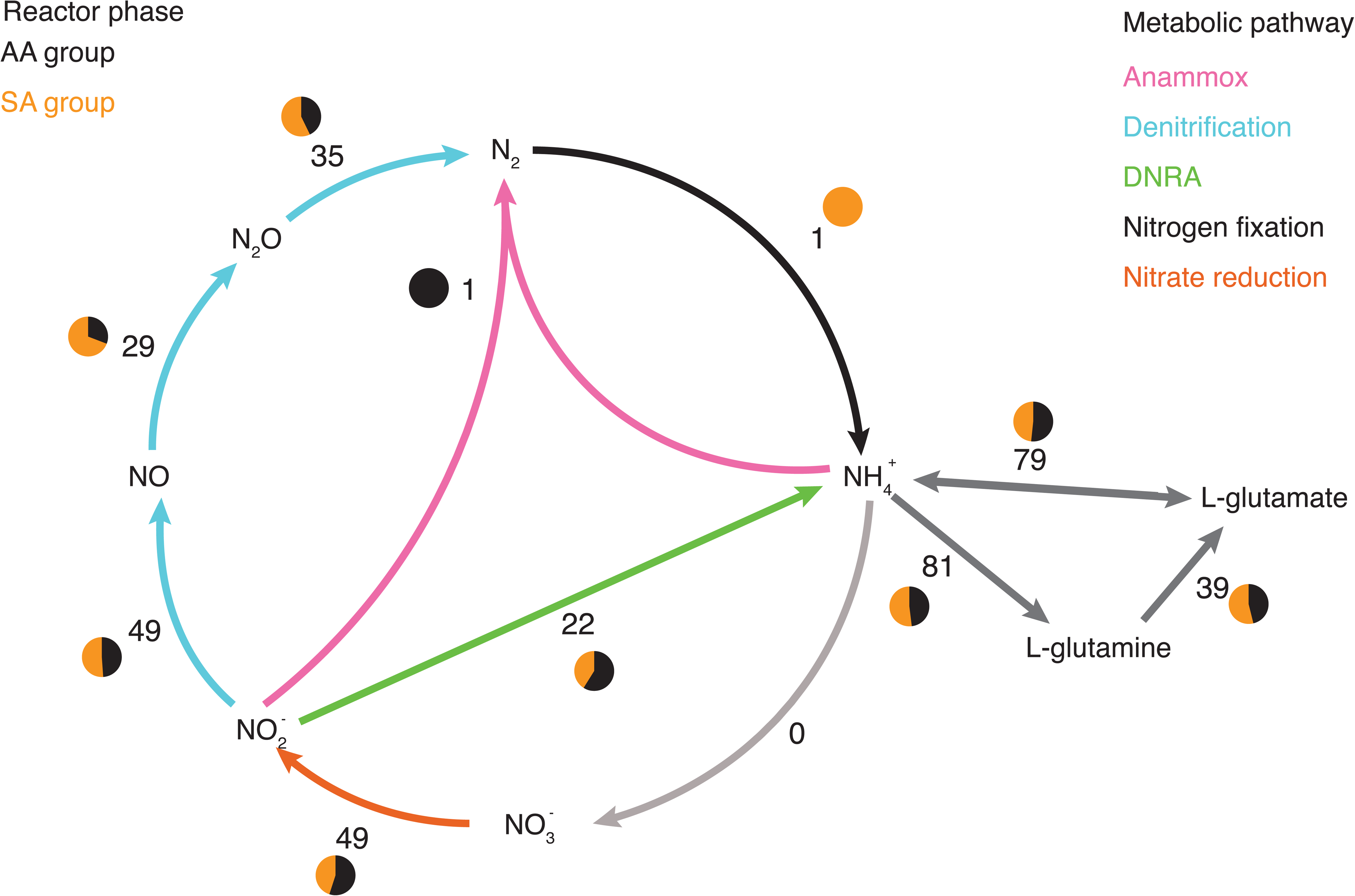
Nitrogen cycle in the anammox bioreactor. The steps in the nitrogen cycle are color-coded by their association to different types of metabolism. The number of bacteria with genes encoding a given step is listed, and the pie chart depicts the ratio between AA and SA bacteria associated with the step.

Carbon fixation is a necessary step in the anammox bioreactor since the influent media did not contain organic carbon. Only two bacteria in the community could be considered autotrophic primary producers. *Brocadia* was confirmed as a primary producer, fixing carbon via the Wood-Ljungdahl pathway and obtaining energy from the anammox pathway. The second bacterium, LAC_PROT27 (*Proteobacteria*, group AA), might be able to fix carbon by the Calvin cycle and obtain energy from denitrification, and could possibly oxidize sulfide to sulfite (*dsrAB* is present in the genome). While LAC_PROT27 was consistently highly abundant in the bioreactor, it was always at least 3-folds less abundant than *Brocadia* (except at time 0). Several other bacteria were also potential autotrophs (or mixotrophs), but were low in relative abundance over the bioreactor’s lifespan. Additional information about carbon metabolism and electron transfer can be found in supplementary information file 1.

### Analysis of metabolic selection in the anammox bioreactor

During the maturation of the anammox bioreactor, the number (Table 1) and diversity (Table S4) of genomes were both reduced. To examine why certain bacteria were enriched (group AA) while others were removed (group SA), we compared the genomes’ ability to synthesize metabolites against their ability to acquire nutrients from the environment. For synthesis we checked 24 KEGG modules for amino acids (aa.), 18 modules for vitamins and cofactors, and 28 modules for lipids and fatty acids. For nutrient acquisition we checked 54 KEGG modules for transporters. The mean module completeness across these categories was compared. A complete module implies a bacterium has the functional capability (be it synthesis or transport). Thus, the higher the module completeness of a group, the more likely it is that its members have the relevant functional capability. For statistical analysis, when both data sets (AA and SA) fit a normal distribution, a two sample T-test was conducted. When the values did not fit a normal distribution, the ratio between groups AA and SA was calculated, and used to set a confidence interval (CI: mean ± 1.64*(SD/n^0.5^), alpha = 0.05). Values outside of the CI were considered significantly different than the mean.

Synthesis modules for aa. (p-value = 0.68) and vitamins/cofactors (p-value = 0.51) both fit a normal distribution, and no statistically significant differences were found for these categories between the AA and SA bacterial groups. Synthesis modules for lipids and fatty acids did not have a normal distribution, so their ratio was inspected (CI upper = 1.22. CI lower = 0.80). Six modules were significantly higher and 14 were significantly lower in the AA group vs. the SA group. Group AA had a higher proportion of Gram (+) bacteria, which also added to the difference in module completeness. Transport modules also did not fit a normal distribution. Inspection of the module ratio showed that both the upper CI (2.69) and lower CI (1.74) were higher than a ratio of 1. Of the 26 transport modules with a ratio >1.74, 18 were transport systems for organic carbon molecules (sugars, lipids, a.a., and cofactors).

These comparisons suggest that a bacterium’s ability to acquire nutrients from its environment may be a selective driver in the anammox bioreactor’s community. This was particularly highlighted when observing Metabolic group α, the dominant metabolic group within the AA bacteria. Members of group α have a cassette of extracellular proteases and decarboxylases paired with a wide array of transporters (Figure 5B) that enable acquisition of nutrients from the environment. In addition, the larger ratio of bacteria with auxotrophies in the AA bacteria (Figure 4A, red rectangles for groups α and δ) hints of a greater reliance on external metabolites from other members of the community.

### Metabolic interdependencies between community members

The bacteria in the AA community have a complex metabolic system, with many bacteria relying on other members to provide them with necessary metabolites. In the mature functioning bioreactor, *Brocadia* was the only primary producer present. It was also the only bacterium capable of synthesizing vitamin B12. For most other metabolites (e.g., vitamins and cofactors) the possible metabolic interdependencies [33] are less straightforward (Figure 7 and Table S5). In the schematic figure, the size of each group reflects its relative abundance in the bioreactor at D437. The arrows point towards the group that potentially receives metabolites it cannot synthesize, and the arrows’ sizes reflect the proportion of the examined metabolites that the group needs. Members of group α (the most dominant group aside from *Brocadia*) had multiple auxotrophies in the synthesis of vitamins, cofactors, fatty acids, and lipids. Group α could produce a few metabolites needed by other groups. Members of group ε were the most metabolically diverse and could produce many metabolites needed by other groups. This group could account for most of the auxotrophies of *Brocadia*. Members of group ε could support 75% of *Brocadia*’s auxotrophies in aa, vitamins and cofactors, as well as 60% of its auxotrophies in fatty acids and lipid synthesis (Table S5). Group γ was the smallest group in the bioreactor at D437 and had a mix of auxotrophies and metabolic support potential.

**Figure 7.**
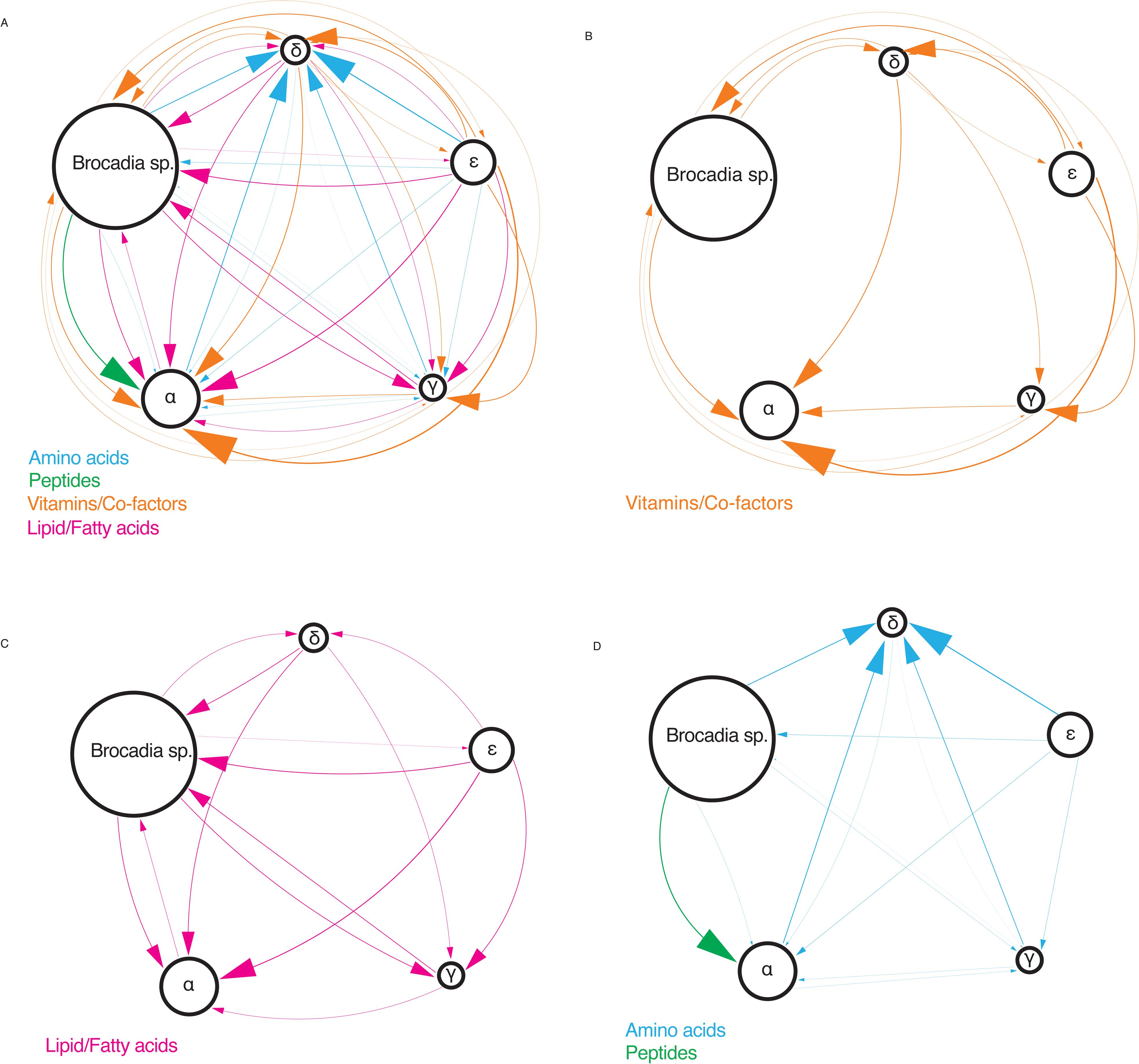
Potential metabolic interdependencies between the metabolic groups in the anammox bioreactor. (A) All potential metabolic interdependencies combined. (B) Metabolic interdependencies for vitamins/cofactors alone. (C) Metabolic interdependencies for lipids/fatty acids alone. (D) Metabolic interdependencies for amino acids and peptides alone. Arrows were assigned according to the absence of a group’s ability to synthesize a metabolite, and they connect to all groups that have the ability (there is redundancy in arrows). The arrowhead points at the group that receives the metabolite. The width of the arrow is proportional to the ratio of metabolites of a given type that are provided; amino acids - 20 metabolites; Peptides - deduced from proteases and transporters (Figure 5B); Vitamins/Co-factors - 10 metabolites; Lipids/Fatty acids - 7 metabolites. The size of each group is proportional to their relative abundance at Day 437. Group β is not shown since it is assumed that all members obtain all of their nutrients and metabolites from their host. Overall, groups □ and δ receive the most metabolites and group ε receives the least. Group δ has the highest number of aa. synthesis auxotrophies and can potentially acquire these from many other community members. Group ε has only a single auxotrophy in vitamin/Co-factor synthesis, while most other groups have multiple auxotrophies. *Brocadia* sp. is the only bacterium capable of vitamin B12 synthesis.

When combining all of the above data, we found that groups □ and ε both had mutualistic associations with *Brocadia* (Figure 7). Group ε potentially provided more metabolites to *Brocadia* than it received, whereas groups □ and δ seemed to gain more from *Brocadia* than they provided.

### Investigating a microbial driven anammox performance destabilization event

Just before day 300 of the bioreactor’s lifespan, an unexpected anammox performance destabilization event occurred. We hypothesized that some interaction between the anammox bacterium and the co-existing community members led to this event. To evaluate which bacteria might have influenced the destabilization event and which might have been influenced, we relied on two parameters: replication and relative abundance (using coverage). First, we checked for changes in a genome’s replication rate (as calculated by iRep [34], see methods for detailed explanation) between D166 and D284. Second, we investigated the log-ratio (LR) of coverage [35] between D284 and D328. To eliminate bias in microbial loading or sequencing depth, we used four different reference frame genomes (RFg) for the LR calculation [35]. Difference in sequencing depth or microbial load can highly bias comparisons between samples. Higher sequencing depth means higher coverage across the entire sample so it might seem that the abundance of bacteria has increased. To remove that bias a reference frame genome(s), with little change in relative abundance over time, is chosen. The abundance values of all other genomes within a given time point are divided by the abundance of the RFg before calculating the log-ratio between samples. The internal ratio removes the aforementioned bias between samples. By combining these two parameters, we were able see which bacteria were actively replicating before the performance destabilization event and eliminate biases created by relative abundance results.

Genomes with iRep values for three out of the four time points between D166 and D437 (27, accounting for > 80% of community) were chosen for this analysis. For each RFg in the LR calculations, significant change was considered for values outside of the confidence interval (CI), calculated for all 127 genomes (Table S6). A bacterium was considered to influence the destabilization if it both increased in replication prior to the event and had a positive and significantly high LR relative to each RFg. A bacterium was considered to be influenced by the destabilization if it decreased its replication rate prior to the event and its LR was significantly low.

Two *Chloroflexi* (LAC_CHLX01 and LAC_CHLX10) showed consistent, significant growth across all RFgs (Figure 8A and Table 2), as well as increased replication rates prior to the destabilization event (Figure 8B). These bacteria likely influenced the destabilization event. Three additional bacteria (the *Chloroflexi* LAC_CHLX06, LAC_CHLX09, and *Ignavibacteria* LAC_IGN05) were also potential influencers, having significant growth based on some of the RFgs. On the other hand, *Brocadia* (LAC_PLT02), showed significantly decreased replication rates prior to the destabilization event. *Brocadia*’s replication rate on D284 (1.07) indicates that only 7% of the bacterium’s population was actively replicating. At other time points, *Brocadia’s* replication rate climbed as high as 2.13, indicating that 100% of the bacterium’s population was actively replicating. One additional bacterium (LAC_PROT22, a *Proteobacteria* from group AA) showed decreased replication and growth (under two RFgs).

**Table 2.** Log-Ratio and replication rates (iRep) for select bacteria.

| Genome | LR<sub>(RF1)</sub> | LR<sub>(RF2)</sub> | LR<sub>(RF3)</sub> | LR<sub>(RF4)</sub> | iRep D166 | iRep D284 |
| --- | --- | --- | --- | --- | --- | --- |
| LAC_CHLX01 | 1.05 | 1.17 | 2.27 | 1.73 | 1.16 | 1.32 |
| LAC_PROT02 | -1.54 | -1.41 | -0.31 | -0.85 | 1.17 | 1.30 |
| LAC_PROT03 | -1.50 | -1.37 | -0.27 | -0.81 | 1.32 | 1.40 |
| LAC_CHLX02 | -2.91 | -2.79 | -1.68 | -2.23 | 1.16 | 1.17 |
| LAC_ACD03 | -0.99 | -0.86 | 0.24 | -0.30 | 1.19 | 1.35 |
| LAC_BACT04 | -0.77 | -0.64 | 0.46 | -0.08 | 1.26 | 1.26 |
| LAC_IGN05 | -0.33 | -0.20 | 0.90 | 0.36 | 1.38 | 1.51 |
| LAC_PROT19 | -2.09 | -1.96 | -0.86 | -1.40 | 1.22 | 1.32 |
| LAC_IGN07 | -0.98 | -0.85 | 0.25 | -0.29 | 1.30 | 1.48 |
| LAC_PROT21 | -1.18 | -1.06 | 0.04 | -0.50 | NA | 1.29 |
| LAC_BAC19 | -0.13 | 0.00 | 1.10 | 0.56 | NA | 1.38 |
| LAC_BAC20 | -1.17 | -1.04 | 0.06 | -0.48 | 1.13 | 1.19 |
| LAC_BACT12 | -1.72 | -1.59 | -0.49 | -1.03 | 1.14 | 1.18 |
| LAC_NIT01 | -1.93 | -1.80 | -0.70 | -1.24 | 1.25 | 1.31 |
| LAC_CHLX06 | -0.18 | -0.05 | 1.05 | 0.51 | 1.21 | 1.28 |
| LAC_PROT22 | -1.23 | -1.10 | 0.00 | -0.54 | 1.36 | 1.24 |
| LAC_PLT02 | -2.12 | -1.99 | -0.89 | -1.43 | 2.13 | 1.07 |
| LAC_CHLX07 | 0.00 | 0.13 | 1.23 | 0.69 | 1.69 | NA |
| LAC_BAC21 | -1.31 | -1.18 | -0.08 | -0.62 | 1.13 | 1.25 |
| LAC_CHLX09 | 0.25 | 0.37 | 1.48 | 0.93 | 1.56 | 1.63 |
| LAC_CHLX10 | 0.43 | 0.55 | 1.66 | 1.11 | 1.26 | 1.42 |
| LAC_BAC22 | 0.27 | 0.40 | 1.50 | 0.96 | NA | 1.39 |
| LAC_BAC23 | -0.31 | -0.18 | 0.92 | 0.38 | 1.17 | 1.17 |
| LAC_PROT27 | -0.35 | -0.23 | 0.88 | 0.33 | 2.06 | 1.56 |
| LAC_PROT28 | -0.69 | -0.56 | 0.54 | 0.00 | 1.33 | 1.27 |
| LAC_PROT30 | -0.48 | -0.36 | 0.75 | 0.20 | NA | 1.89 |
| LAC_D-T02 | -0.43 | -0.30 | 0.80 | 0.25 | NA | 1.59 |

**Figure 8.**
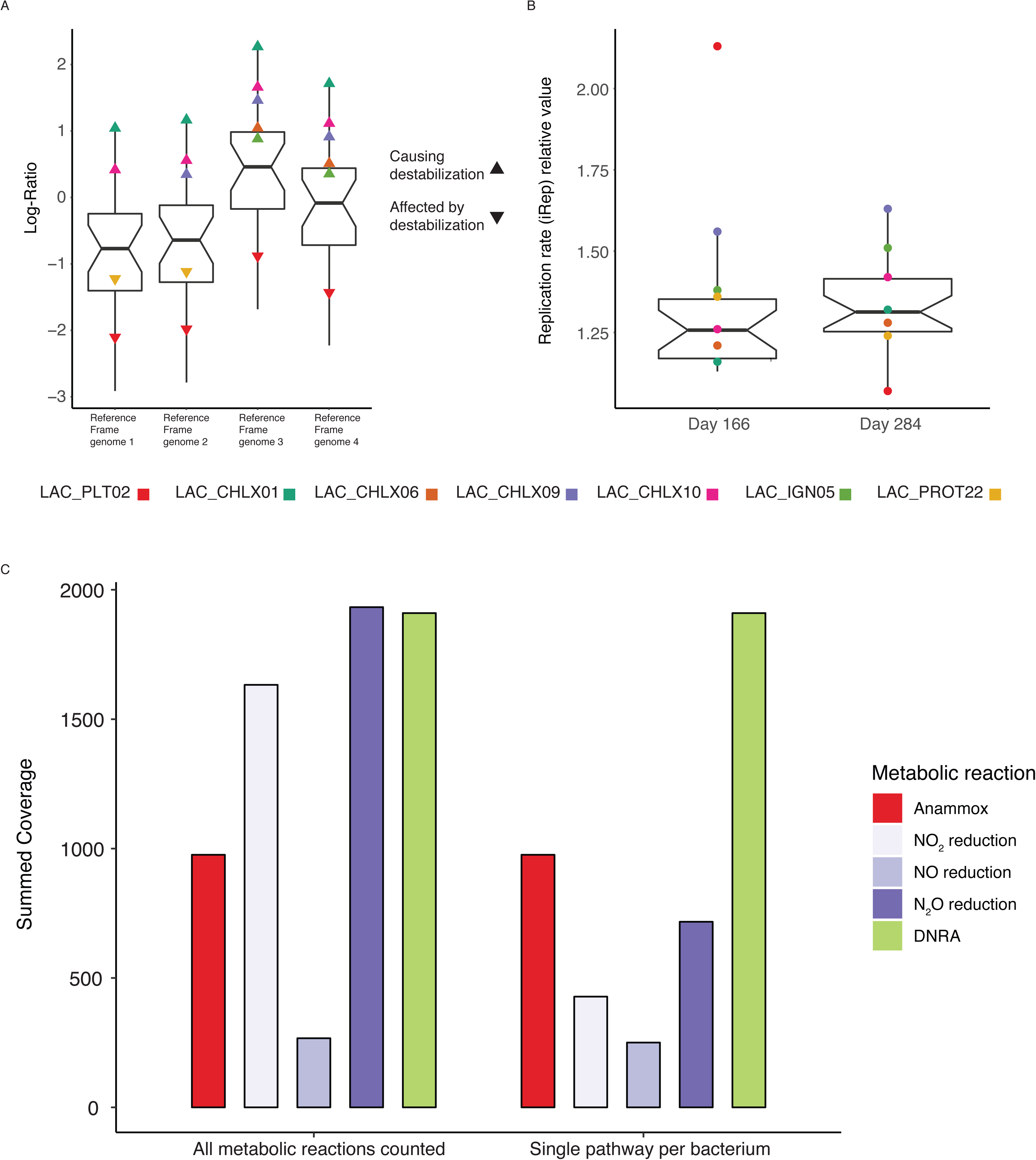
Tracking bacterial growth and nitrogen metabolism in relation to the anammox performance destabilization. (A) Distribution of Log-Ratio changes for selected bacteria between D328 and D284, using different genomes as reference frames. Bacteria that are significantly affected or considered to affect the destabilization event are color-coded. Down-pointing triangles depict bacteria negatively affected by the destabilization and upwards-pointing triangles indicate bacteria that likely caused (or contributed) to the destabilization. (B) Replication rate values on days 166 and 284. Bacteria are color coded the same as in panel A. (C) Relative abundance of nitrogen metabolism pathways in the selected bacteria. Denitrification is divided into its reaction steps. Anammox is considered as a single path since only a single bacterium can perform it (*Brocadia* – LAC_PLT02). DNRA from NO2 is a single step reaction. Abundance was calculated twice, once allowing multiple pathways to occur within each bacterium, and once after choosing a single pathway per bacterium (based on potential energy gain).

Next, we next investigated the nitrogen metabolism of the 27 bacteria, specifically the three metabolic paths that compete for nitrite i.e., anammox, denitrification, and DNRA (Figure 8C). Denitrification was subdivided into three steps (NO_2_ reduction, NO reduction, and N_2_O reduction); DNRA is a single step process. Anammox could only be performed by *Brocadia*. Only a single bacterium (LAC_BAC20) could perform full denitrification. All bacteria capable of DNRA were also capable of partial denitrification. When allowing for a bacterium to have multiple pathways (Figure 8C), NO_2_ reduction via DNRA (*nrfAH*) and denitrification (*nirS/nirK*) appeared to be equally dominant during the anammox performance destabilization event (D328). While N_2_O reduction was also dominant, a bottleneck appeared to exist at the second step, NO reduction. When we assume that the bacteria capable of both DNRA and partial denitrification choose the path that yields the most energy [36], we can remove the partial denitrification pathway (Figure 8C). Under this assumption, DNRA was clearly revealed as the dominant process occurring in the bioreactor during the anammox performance destabilization event. All bacteria that were shown to affect the destabilization event were DNRA bacteria. In addition, the destabilizing bacteria belong to metabolic groups α (LAC_CHLX01, LAC_CHLX06, LAC_CHLX09, and LAC_CHLX10) and δ (LAC_IGN05). Both of these groups were reliant on other members for organic carbon (Figure 5B and Figure 7).

### Core anammox community

Resulting genomes from our study, in combination with genomes from two previous anammox metagenomic studies, Speth et al. [18] (22 genomes) and Lawson et al. [20] (15 genomes), provide strong evidence to support a core anammox community (Figure 9). The relative abundances of bacteria from the dominant phyla across these three bioreactors were fairly similar: in each bioreactor the anammox, along with *Chloroflexi*, *Ignavibacteria*, and *Proteobacteria* bacteria, composed >70% of the community (Figure 9B).

**Figure 9.**
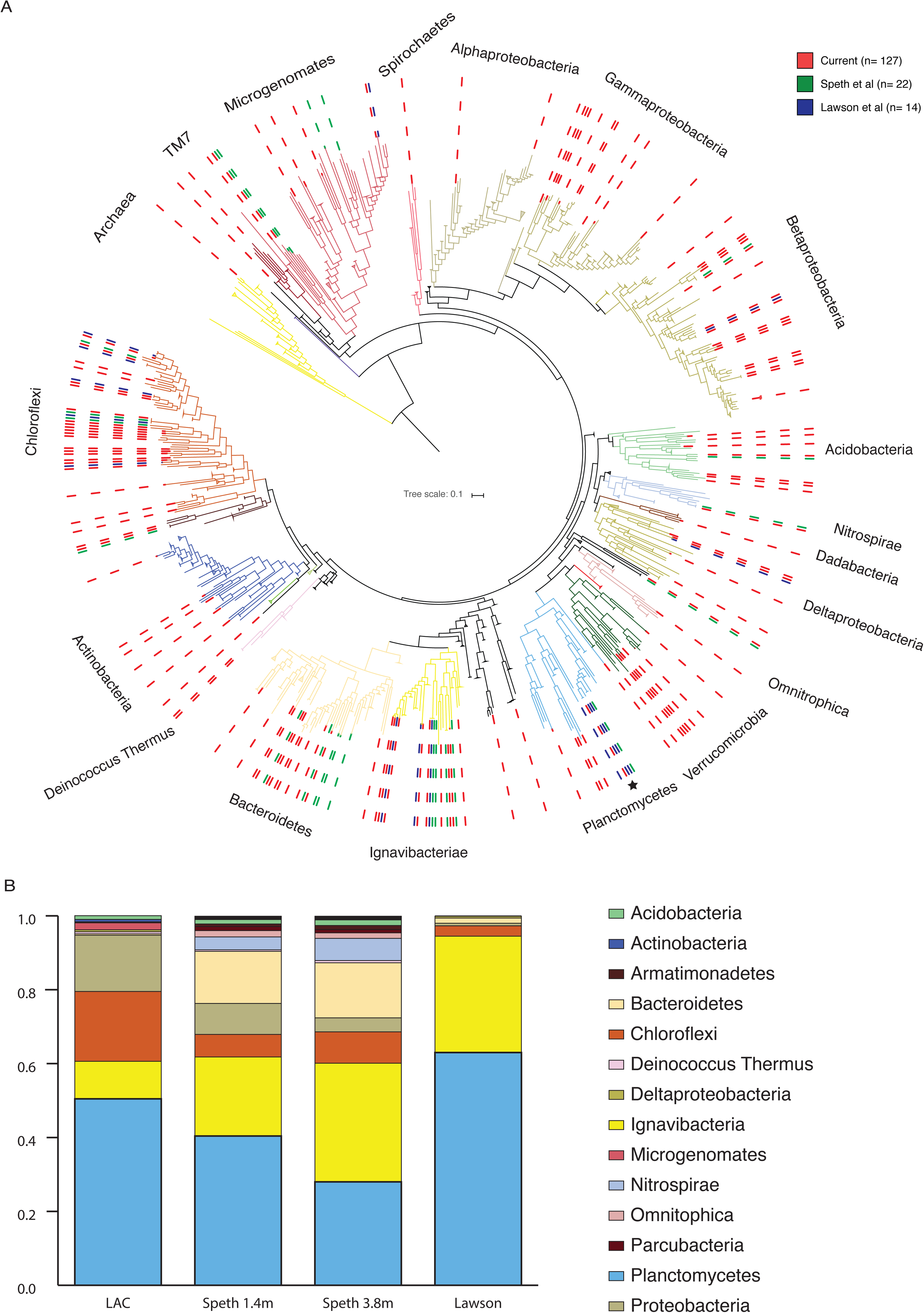
Phylogenetic analysis of three anammox microbial communities. (A) A maximum likelihood tree based on the alignment of 15 concatenated ribosomal proteins. In the construction of the tree 3225 reference sequences were used, with genomes from current and previous genome-centric studies on anammox communities. Genomes from the current anammox community are marked with a red dashed line, genomes from two previously studied communities; Speth et al. and Lawson et al., are marked with green and blue dashed lines, respectively. Nodes containing only reference genomes were collapsed for ease of view. Collapsed nodes are depicted as triangles and their size is relative to the number of bacteria they contain. A black star marks *Brocadia*. (B) relative abundance of major phyla in the three microbial communities. Current community reference data was calculated from day 437 only. The relative abundance of *Brocadia* sp. comprises nearly all of the relative abundance attributed to phylum *Planctomycetes* (with small contributions from other members of the phylum). The most abundant phyla (*Chloroflexi*, *Ignavibacteria*, and *Proteobacteria*) consistently account for >70% of the communities. The phyla colors follow the ggkbase color scheme and the major phyla are shown in the legend.

Due to the significantly larger genome yield and time-series analysis in this study, our bioreactor shared more genomes with each of the other bioreactors than the other bioreactors shared between themselves. In total, 21 genomes from our bioreactor were closely related to those from at least one of the two other bioreactors, 17 of which were present at the last time point, D437 (Table S7). The related bacteria accounted for 50% and 93% of the Speth et al. and Lawson et al. genomes, respectively. The bioreactor studied by Speth et al. was different from the other two bioreactors because it was amended with oxygen to perform partial nitritation and anammox within the same bioreactor, while the others performed anammox only.

A more focused phylogenetic tree of *Planctomycetes* shows that the *Brocadia* in our bioreactor and in the Lawson et al. bioreactor are the same species (*Brocadia sapporensis* [27]), while the *Brocadia* species from the Speth et al. bioreactor is different (*Brocadia sinica*) (Figure S3).

## Discussion

In this study we present an in-depth analysis of the development of an anammox community from seed to stable state (through several perturbations) in an anaerobic membrane bioreactor. By combining several methodologies, we are able to gain important insights into the dynamics and interactions of more than 100 species in the bioreactor community [37, 38].

The first perturbation of the bioreactor, a mechanical malfunction combined with inoculum amendments, changed the trajectory of the community succession. This can be seen from the relative abundance-based grouping (Figures 2-3) and the detected strain shifts. The first inoculum amendment had a much stronger effect on community assembly than the subsequent bioreactor malfunction and second amendment [39, 40]. The large shift in the community occurred between days 96 and 152, after which the community trajectory became fairly consistent until day 290. The oscillations after days 350 are likely due to differences in sequencing depths. Large changes in community structure due to inoculations are a real concern for large-scale bioreactors, where the influent contains constantly-changing communities of bacteria [41, 42]. It is unclear if members of group C, which were dominant in the nascent anammox community, would have better supported the bioreactor’s anammox performance [39]. Looking at their metabolism may offer a few clues: of the 14 genomes clustered into the nascent anammox community, six were of group ε and four were of group γ, both of which were considered to maintain a mutualistic or commensal relationship with the anammox bacterium. In addition, denitrification was more common than DNRA in group C. Two genomes (LAC_ACD03 and LAC_PROT30) from the nascent anammox community were included in the focused investigation of the anammox performance destabilization. Both had no significant effect on the event nor were they significantly affected by it.

The metabolic analysis of the matured community showed that transport systems for nutrients (mainly organic carbon) were the most enriched in the community. The ability of bacteria to utilize available nutrients in the environment [43] has been shown before and has been proposed to explain dominance in nutrient-rich environments [44]. However, when such bacteria favor acquisition over synthesis, it can further stress a community that is reliant on a slow growing primary producer for nutrients [45–48]. Members of groups α and δ have these characteristics, and some are implicated in the anammox performance destabilization event. The aforementioned destabilization event took place after nearly 100 days of high performance, and no external factors could explain the sudden destabilization and performance crash. By combining information about replication rates and changes in relative abundance of community members, we were able to identify several bacteria that likely affected the performance crash. Analysis of composite data (such as relative abundance) for true changes in community structure has many pitfalls [35]. In addition, the response of a bacterium to an event, in this case an increase in relative abundance following the performance destabilization, cannot be deduced to mean it had any effect on the event. We were fortunate to have taken a metagenomic sample the week prior to the destabilization event. By measuring replication rate changes as well as relative abundance changes, we could better deduce a possible causative effect. A bacterium that increases its replication rates prior to the event and increases in relative abundance due to the event is more likely to have a causative effect. While most bacteria had higher replication rates at D284 compared to D166 (17 of 22 bacteria with values in both days), only five bacteria significantly increased in relative abundance following the event. Genes conferring DNRA and partial denitrification capabilities were detected in these bacteria. These types of bacteria could improve bioreactor performance if they remove nitrate and excess nitrite, but they could be detrimental if they compete with anammox for nitrite or allow for the build-up of nitrite. Here, the equilibrium between support of the anammox process and disruption of the anammox process was tilted towards the latter.

Two possible scenarios for nitrogen metabolism are consistent with the bioreactor’s performance that exhibited decreased nitrogen removal and increased ammonium in the effluent leading up to the destabilization event. One scenario is described by DNRA dominance and the second by nitrite reduction to nitric oxide and its subsequent leakage from the system. The first scenario provides more energy to the bacteria [36], so we speculate it is more likely. *Brocadia* has genes required for DNRA, but given the high nitrogen removal rate by the anammox process leading up the disturbance, it can be assumed that *Brocadia* would not be primed to carry out the DNRA reaction on such a short timescale. However, DNRA could potentially be used by *Brocadia* for detoxification by cycling potentially toxic excess nitrite back to ammonium, where it could then participate in the anammox reactions [18, 20].

The replication rate of *Brocadia* prior to the performance destabilization was down to 1.07, from 2.13 on D166. A replication rate of 1.07 is equivalent to only 7% of the population actively replicating, which can explain the large decrease in relative abundance in the following time point. This also hints that the process leading to the destabilization event occurred prior to D284. Whether the process could be assigned to a single specific occurrence, or if it is an additive process culminating in a break from equilibrium among members of the community, is a gap that future research should address.

A broader investigation of metabolic interdependencies within the community shed light on the stability of the anammox community. *Brocadia* is the source of organic material in the community, but obtains essential metabolites from community members, especially *Proteobacteria*. This forms a basis for a mutual symbiotic relationship. On the other hand, *Chloroflexi*, comprising the largest group of bacteria besides *Brocadia*, receive numerous metabolites while apparently providing few in return. They are characterized by an array of extracellular proteases and amylases, likely used to breakdown the extracellular matrices formed by *Brocadia*. *Chloroflexi*, as a group, are most associated with anammox bacteria and form a large fraction of the core community. They also account for the majority of the destabilizing bacteria. Together, the results point to a parasitic symbiosis. While anammox bacteria generate sufficient organic carbon to support the growth of its co-occurring heterotrophic microorganisms, the tipping point between stable and unstable operation and the factors that control it have not been fully identified. Input changes may be able to restore anammox activity, but this is only an empirical solution. In full-scale anammox bioreactors where influent organic carbon is essentially ubiquitous, heterotrophic dominance could persist without some sort of active countermeasure. Therefore, future research should target the inhibition of potential destabilizing heterotrophs.

Previous studies have discussed a potential core anammox community [12–16]. With the exception of very few studies, all such work has been conducted with single gene markers. Our analysis of an anammox community is the largest to-date and thus expands the ability to test this hypothesis. Our results support the existence of a core community, while identifying factors that differentiate communities. The high similarity among bacterial communities originating from three distinct anammox bioreactors [18, 20] strongly suggests a global core anammox microbial community. In the construction of the phylogenetic tree, we used >3000 reference genomes originating from diverse environments. Through this analysis, we found that the anammox community forms distinct clades at the species level, despite the sheer number and diversity of sources. More than half of the bacteria did not have species level relatives, and an additional 26% only had a relative found in our anammox bioreactor or in a previous anammox study [18, 20]. Together, nearly 80% of the bacteria are unique to anammox bioreactors, so it is clear that the anammox bioreactor selects for a unique set of bacteria. Parameters that increased the differences between communities are the species of the anammox bacterium and the bioreactor configuration. Since both parameters relate to the same bioreactor [18], we cannot conclude which has a stronger effect.

## Conclusions

Here we present the largest-to-date metagenomic analysis of a microbial community in an anammox bioreactor. Our results support the growing body of literature that suggests that anammox communities are unique and may share a core microbial community. We identified a distinct phylogenetic profile across reported metagenomic analyses of anammox bioreactors. In subsequent analyses of our metagenomes, we identified metabolic traits associated with the core microbial community that are distinguishable from other bacteria present in the source sludge inoculum. In addition, our time-series analysis included a biologically-driven period of anammox performance destabilization. We identified an increase in replication rates for several bacteria just prior to the event. Further analysis revealed that these bacteria contain genes conferring DNRA, which puts them in direct competition with the *Brocadia* sp. for nitrogen resources. Combined, our results provide a possible mechanistic explanation for the performance shift of the anammox bioreactor and advance the comprehensive control of this promising technology. However, further work is needed to elucidate the precise mechanisms that govern anammox community interactions and to predict performance destabilization events.

## Methods

### Bioreactor operation

A laboratory-scale, anammox anaerobic membrane bioreactor (MBR) with a working volume of 1L was constructed and operated for over 440 days (Figure S4). The bioreactor was originally inoculated with approximately 2 g volatile suspended solids (VSS) L^-1^ of biomass from a pilot-scale deammonification process treating sidestream effluent at San Francisco Public Utilities Commission (SFPUC) in San Francisco, CA. The bioreactor was re-inoculated with similar concentrations of biomass from the same source on Days 147 and 203. Synthetic media containing ammonium, nitrite, bicarbonate, and trace nutrients (meant to mimic sidestream effluent at a municipal wastewater treatment plant) was fed to the bioreactor (Table S8). For the first 154 days of operation, the bioreactor was kept under nitrite-limiting conditions to prevent inhibitory conditions due to the buildup of nitrite, and influent ammonium and nitrite concentrations ranged from 200-300 mg N L^-1^ and 100-300 mg N L^-1^, respectively. On Day 154, ammonium and nitrite concentrations were adjusted to the theoretical anammox stoichiometric ratio, 1:1.32. Afterwards, influent ammonium and nitrite concentrations were maintained at this ratio. Ammonium ranged from 200-500 mg N L^-1^ and nitrite 265-660 mg N L^-1^. On Day 353, influent concentrations of copper, iron, molybdenum, and zinc were increased based on literature suggestions [22–25].

The bioreactor was operated in a continuous flow mode. For the first 145 days, the hydraulic retention time (HRT) was maintained at 48 hours; afterwards it was reduced to 12 hours. No solids were removed from the bioreactor for the first 100 days of operation; afterwards, the solids retention time (SRT) was reduced to 50 days. A polyvinylidene fluoride hollow fiber membrane module with a 0.4 µm pore size and total surface area of 260 cm^2^ (Litree Company, China) was mounted in the bioreactor. Temperature was maintained at 37° C with an electric heating blanket (Eppendorf, Hauppauge, NY). Mixing was provided by an impeller at a rate of 200 rpm. Mixed gas was supplied continuously to the bioreactor (Ar:CO_2_ = 95:5; 50 mL min^-1^) to eliminate dissolved oxygen and maintain a circumneutral pH range of 6.9-7.2. Influent and effluent concentrations of ammonium, nitrite, and nitrate were measured approximately every other day using HACH test kits (HACH, Loveland, CO), as described in the manufacturer’s methods 10031, 10019, and 10020, respectively.

### Biomass collection and DNA extraction

Biomass samples were extracted via syringe from the bioreactor every 2-10 days, flash frozen in liquid nitrogen, and stored frozen at −80 °C until use. Genomic DNA was extracted from the samples using the DNeasy PowerSoil Kit (Qiagen, Carlsbad, CA), as described in the manufacturer’s protocol. Extracted DNA was quantified with a NanoDrop Spectrophotometer (Thermo Scientific, Waltham, MA), and normalized to approximately 10 ng/µL with nuclease-free water (Thermo Scientific, Waltham, MA). All genomic DNA samples were stored at −20 °C until use. For shotgun metagenomic sequencing, samples were sent to the Joint Genome Institute (JGI) in Walnut Creek, CA. There, DNA quality was assessed prior to library preparation and sequencing (150bp pair-end) on the Illumina HiSeq 2500 1T sequencer (Illumina, San Diego, CA). For 16S rRNA sequencing, samples were sent to the Institute for Environmental Genomics at the University of Oklahoma. There, DNA quality was assessed prior to library preparation and amplicon sequencing on the Illumina MiSeq sequencer (Illumina, San Diego, CA).

### Metagenomic sequencing, assembly, and binning

Resulting sequences from each time point were processed separately, following the ggKbase SOP (https://ggkbase-help.berkeley.edu/overview/data-preparation-metagenome/). Briefly, Illumina adapters and trace contaminants were removed (BBTools, GJI) and raw sequences were quality-trimmed with Sickle [49]. Paired-end reads were assembled using IDBA_UD with the pre-correction option and default settings [50]. For coverage calculations, reads were mapped with bowtie2 [51]. Genes were predicted by Prodigal [52] and predicted protein sequences were annotated using usearch [53] against KEGG, UniRef100, and UniProt databases. The 16S rRNA gene and tRNA prediction was done with an in-house script and tRNAscanSE [54] respectively. At this point, the processed data was uploaded to ggKbase for binning.

Manual binning was performed using the ggKbase tool. The binning parameters were GC% and coverage (CV) distribution, and phylogeny of the scaffolds. Quality of the manual bins was assessed by the number of Bacterial Single Copy Genes (BSCG) and Ribosomal Proteins (RP) found in each bin (aiming at finding the full set of genes, while minimizing multiple copies). In addition to manual binning, automated binning was done using four binners: ABAWACA1 [30], ABAWACA2, CONCOCT [55], and Maxbin2 [56]. For all, the default parameters were chosen.

All bins from both automatic and manual binning tools were input into DASTool [57] to iterate through bins from all binning tools and choose the optimal set of bins. checkM was run to analyze genome completeness [26]. The scaffold-to-bin file created by DASTool was uploaded back to ggKbase and all scaffolds were rebinned to match the DASTool output. Each of the new bins were manually inspected and scaffolds suspected of being falsely binned were removed.

After inspecting the first round bins, we improved the high coverage bins by subsampling the read files, and repeating the SOP above [58]. In addition, refinement of the *Brocadia* Genome bins was done with ESOMs [59] (Supplemental methods).

### Post binning analysis

Unique representative genomes were determined by the dereplication tool, dRep [60], using a 95% threshold for species level clustering. Within each cluster, the representative genome was chosen based on their completeness, length, N50, contamination, and strain heterogeneity. In several clusters with higher heterogeneity, a second strain was chosen (Table S2). The strain threshold was set at 2% difference.

All representative and strain genomes were curated by correcting scaffolding errors introduced by idba_ud, using the ra2.py program [30]. Following curation, the genomes were processed again for gene calling and annotation (see above for details). The replication rates of bacteria can be inferred from examining the coverage ratio between the origin of replication and the terminus of replication. In a population that is not replicating there will be no difference in the coverage so the ratio would be one. If the population is replicating, we expect the ratio to be >1 since there would be replication forks that have not finished replicating and thus the coverage towards the origin of replication would be higher than that of the terminus. Calculating replication rate is more complicated in metagenomic samples but it is still possible to look at overall trends in coverage across the genome. Analysis of replication rates at different time points was performed with the iRep program [34] using the default parameters (Table S9). Briefly, iRep calculates replication rate by measuring sequencing coverage trend that results from bi-directional genome replication from a single origin of replication. The program uses only high-quality draft genomes (>=75% complete, <=175 fragments/Mbp sequence, and <=2% contamination). Since iRep is a measure of a trend it does not have any units.

Raw reads were submitted to the National Center for Biotechnology Information (NCBI) Genbank, under project accession number PRJNA511011. In addition, the Representative and Strain genomes were uploaded to ggkbase as two separate projects (https://ggkbase.berkeley.edu/LAC_reactor_startup/organisms and https://ggkbase.berkeley.edu/LAC_reactor_strains/organisms).

### Phylogenetic analysis and core anammox analysis

The taxonomic affiliation of each genome was initially assigned in ggKbase, based on the taxonomic annotation of genes in the scaffolds. For each hierarchical taxonomic level, the taxonomy was decided if at least 50% of genes had a known taxonomic identification.

Phylogenetic analysis of the genomes (current study, Speth et al. [18], and Lawson et al. [20]) was based on a set of 15 ribosomal proteins [61]. Each gene was aligned separately to a set of 3225 reference genomes, followed by concatenation while keeping the aligned length of each gene intact. A preliminary tree was created by adding the queried genomes to the reference tree using pplacer v1.1.alpha19 [62] and a set of in-house scripts. The tree was uploaded to iTOL [63] for visualization and editing. After initial inspection we decided to reduce the tree in preparation for creating a maximum likelihood tree. Large phyla with no representatives in an anammox sample were removed (approximately 1000 sequences). The remaining sequences were aligned by MUSCLE [64] and a RAxML tree built in The CIPRES Science Gateway V. 3.3 [64, 65].

For the analysis of phylogenetic distance between different anammox community members, we used the APE package [66] in R [67, 68] to extract the distance matrix. Species level distance was set at 5% of the longest measured distance on the tree. The R script and RData files for the analysis of related species, community dynamics, and metabolic capacities were uploaded to figshare (https://figshare.com/projects/Anammox_Bioreactor/59324).

### 16S rRNA gene sequencing, processing, and analysis

DNA samples, taken at 55 time points across the lifespan of the bioreactor, were sent to the Institute for Environmental Genomics at the University of Oklahoma (Norman, OK) for amplification of the variable 4 (V4) region of the 16S rRNA gene, library preparation, and amplicon sequencing. The full protocol was previously described in Wu et al. [69]. In summary, the V4 region of the bacterial 16S rRNA gene was amplified from DNA samples using primers 515F (5’-GTGCCAGCMGCCGCGG-3’) and 806R (3’-TAATCTWTGGVHCATCAG-5’), with barcodes attached to the reverse primer. Amplicons were pooled at equal molarity and purified with the QIAquick Gel Extraction Kit (QIAGEN Sciences, Germantown, MD). Paired-end sequencing was then performed on the barcoded, purified amplicons with the Illumina MiSeq sequencer (Illumina, San Diego, CA).

Subsequent sequence processing and data analysis were performed in-house using MOTHUR v.1.39.5, following the MiSeq SOP [70, 71]. In summary, sequences were demultiplexed, merged, trimmed, and quality filtered. Unique sequences were aligned against the SILVA 16S rRNA gene reference alignment database [72]. Sequences that did not align to the position of the forward primer were discarded. Chimeras were detected and removed. The remaining sequences were clustered into operational taxonomic units (OTUs) within a 97% similarity threshold using the Phylip-formatted distance matrix. Representative sequences from each OTU were assigned taxonomic identities from the SILVA gene reference alignment database [72]. Sequences that were not classified as bacteria were removed. Remaining OTUs were counted, and the 137 most abundant OTUs (accounting for up to 99% of sequence reads within individual samples) were transferred to Microsoft Excel (Microsoft Office Professional Plus 2016) for downstream interpretation and visualization. The read files from all time points, as well as the 137 most abundant OTUs were uploaded to figshare (https://figshare.com/projects/Anammox_Bioreactor/59324).

In order to correlate genome-based OTUs to 16S rRNA gene-based OTUs, 16S rRNA gene sequences were extracted from the representative genomes and combined with the representative sequences from the 137 most abundant 16S rRNA gene-based OTUs. If a representative genome did not contain the V4 region of the 16S rRNA gene, the region was pulled from another genome in the same cluster. The combined 16S rRNA gene sequences were aligned following the protocol described above, and those sharing at least 99% average nucleotide identity were assumed to represent the same microorganism [73].

### Community dynamics analysis

The paired sequence reads from all time points were mapped to the set of reference genomes using bowtie2 [51], followed by calculations for coverage (average number of reads mapped per nucleotide) and breadth (percent of genome covered by at least one read) for each genome per time point [74]. The multiplication of the two values was then used to calculate the estimated relative abundance. These steps were done to negate the biases created by repetitive sequences that occur more often in partial genome bins.

Association between genomes was tested by calculating pairwise correlation for all genomes by relative abundance. Rho values (ranging from −1 to 1) were used to create a distance table (Euclidean distance), followed by clustering with the ward.D method. The resulting clusters were marked A-D. To test for the association of genomes and clusters to time points, we ran a nMDS analysis (non-parametric MultiDimensional Scaling) with genomes and time points. Each genome was colored by its relative abundance cluster on the 2D projection of the nMDS.

For relative abundance changes, the estimated relative abundances of genomes were divided by the sum of all estimated relative abundance values per time point. For a clearer resolution of changes in the four relative abundance groups, the *Brocadia* (part of group D) was presented separately.

### Metabolic analysis

The functional profiles of the genomes were evaluated using KEGG KAAS [75], with Hidden Markov Models for shared KEGG orthologies (KOs) [28, 29, 76]. From this, we received the KEGG annotation (KO number) for all open reading frames and a completeness value for each KEGG module. KO annotations that were questionable were removed from the analysis.

From the KO list, we created a presence-absence matrix (Jaccard index), and clustered the genomes using the Complete method. From module completeness, we created a Euclidean distance matrix, followed by clustering with the ward.D method. Based on module completeness clustering, we assigned genomes to metabolic groups □-ε.

For each metabolic group, a representative metabolic map was created. Module completeness greater than 67% in at least half of the group members was considered representative of the group. Once the modules were selected, they were drawn and connected based on metabolic KEGG maps. Additional reaction, complexes, and transporters were added according to KO presence (e.g., aa. synthesis, oxidative phosphorylation complexes, flagellar motor, etc.).

For nitrogen metabolism, all relevant KOs were examined. For the purpose of this study, nitrate reduction was considered a separate path from denitrification/DNRA, since it could be the first step in both pathways, using the same enzymes. Denitrifying bacteria were considered to be bacteria capable of full conversion of nitrite to N_2_. DNRA bacteria were considered to be bacteria capable of conversion of nitrite to ammonium using the nrfAH enzymes. No partial nitrogen process was considered for this paper, although it was present, according to per-step analysis.

## Notes

Ray Keren and Jennifer E. Lawrence share senior authorship.

## List of abbreviations

Anammox: anaerobic ammonium oxidation
DNRA: dissimilatory nitrate reduction to ammonium
PN: partial nitritation
NRR: nitrogen removal rate
D0: day 0
D82: day 82
D166: day 166
D284: day 284
D328: day 328
D437: day 437
ANI: average nucleotide identity
nMDS: nonmetric multidimensional scaling
AA: anammox associated
SA: source Associated
HMM: Hidden Markov Model
KEGG: Kyoto Encyclopedia of Genes and Genomes
KO: KEGG Orthology
CPR: Candidate Phyla Radiation
CI: confidence interval
LR: log-ratio
RFg: reference frame genome.

## Declarations

### Ethics approval and consent to participate

Not applicable

### Consent for publication

Not applicable

### Availability of data and materials

The metagenomic datasets analyzed during the current study are available on the National Center for Biotechnology Information (NCBI) Genbank, under project accession number PRJNA511011.

The 16S rRNA gene datasets analyzed during the current study are available on the Figshare repository: https://figshare.com/projects/Anammox_Bioreactor/59324

The R script used to analyze the metagenomic data is also available on the Figshare repository, under the same link as above.

All other data generated during this study is either included in this published article (and its supplementary information files) or can be made available from the corresponding author upon reasonable request.

### Author contribution

K.Y. and L.Z. supervised the study. K.Y., L.Z., L.A-C., and D.J. designed the study. L.Z. built and maintained the bioreactor. R.K. analyzed metagenomics data and wrote the manuscript. J.L. analyzed 16S rRNA gene data, analyzed bioreactor performance, and wrote the manuscript. J.F.B. supervised the metagenomics analysis. W.Z. and K.Y. contributed to bioreactor maintenance and analysis, sampling, and 16S rRNA gene data analysis. All authors read the manuscript and contributed with their input.

## Acknowledgements

The authors would like to thank Edmund Antell for his help with evaluating energy gains via nitrogen metabolism, as well as his feedback on manuscript revisions.

## Additional Files

Additional file 1: **Table S1** Genome parameters for the representative bacteria from the anammox bioreactor. **Temporal strain shifts in the bioreactor**. Analysis of shifts between bacterial strains during the first bioreactor crash and subsequent biomass amendment. **Table S2.** Genome parameters for the bacterial strains related to representative bacteria from the bioreactor. **Table S3** assignment of genomes to different clusters. **Figure S1.** Analysis of community metabolism by gene presence. **Figure S2.** Reduced metabolic map of the Brocadia sp. Bacterium. **Electron transfer pathways in the community.** Analysis of different electron donors and acceptors utilized by the anammox community members. **Central carbon metabolism in the community**. Analysis of carbon utilization in the anammox microbial community. **Table S4** Diversity indices of the anammox community. **Table S5**. Synthesis and transport of metabolites in the metabolic groups. **Table S6** Confidence intervals of the Log-Ratio changes for each Reference Frame genome. **Table S7.** Closest relative based on distance in ML tree for anammox bacterial community. **Figure S3.** Maximum likelihood tree of the PVC clade (Planctomycetes, Verrucomicrobia, Chlamydiae, and Lentisphaerae). **Figure S4.** Schematic of the anaerobic membrane bioreactor. **Table S8**. Composition of the synthetic wastewater feed. **Supplemental methods.** Refinement of the *Candidatus Brocadia* sp. genome. **Table S9**. Replication rates of genomes for each time point, calculated using the iRep program.

