## supplemental information for "Increased Replication of Dissimilatory Nitrate-Reducing Bacteria Leads to Decreased Anammox Bioreactor Performance"

**Affiliations**
1- Department of Civil and Environmental Engineering, University of California Berkeley, Berkeley, CA, USA
2- Tighe & Bond, Westwood, MA, USA
3- Department of Civil and Environmental Engineering, The University of Auckland, Auckland, New Zealand
4- Department of Earth and Planetary Sciences, University of California Berkeley, Berkeley, CA, USA
5- Earth and Environmental Sciences Division, Lawrence Berkeley National Laboratory, Berkeley, CA, USA
6- College of Chemistry and Environmental Engineering, Shenzhen University, Shenzhen, China
7- Shenzhen Graduate School, Peking University, Shenzhen, China

**Table S1** Genome parameters for the representative bacteria from the anammox bioreactor

| Genome | Genome code (ggkbase) | Phyla | Genome length (bp) | GC% | Contigs | ORFs | Completeness (%) |
|---|---|---|---|---|---|---|---|
| LAC_ACD01 | anamox1_Acidimicrobiia_62_4_curated | *Acidobacteria* | 298722 | 58.08 | 173 | 496 | 65.54 |
| LAC_ACD02 | anamox1_Acidobacteria_62_5_curated | *Acidobacteria* | 3111074 | 61.59 | 565 | 3251 | 65.54 |
| LAC_ACD03 | anamox1_Pyrinomonas_methylaliphatogenes_54_8_curated | *Acidobacteria* | 2071852 | 54.23 | 87 | 1989 | 95.69 |
| LAC_ACD04 | anamox3_Acidobacteria_71_4_curated | *Acidobacteria* | 1334157 | 69.62 | 772 | 1759 | 39.71 |
| LAC_ACD05 | LAC_NA06_Candidatus_Solibacter_usitatus_62_12_curated | *Acidobacteria* | 4513182 | 61.85 | 167 | 4021 | 91.38 |
| LAC_ACD06 | LAC_NA07_Acidobacteria_70_38_curated | *Acidobacteria* | 3262818 | 69.91 | 40 | 2917 | 94.83 |
| LAC_ACD07 | LAC_NA07_Candidatus_Solibacter_usitatus_59_6_curated | *Acidobacteria* | 1779371 | 57.94 | 742 | 2242 | 56.02 |
| LAC_ACT01 | anamox2_Actinobacteria_65_5_curated | *Actinobacteria* | 2799571 | 63.61 | 662 | 3147 | 67.46 |
| LAC_ACT02 | LAC_NA07_Actinobacteria_71_14_curated | *Actinobacteria* | 3047960 | 70.65 | 64 | 3117 | 93.97 |
| LAC_ACT03 | LAC_NA07_Actinobacteria_74_18_curated | *Actinobacteria* | 1235923 | 73.53 | 141 | 1261 | 64.86 |
| LAC_ACT04 | LAC_NA07_Actinotalea_fermentans_75_19_curated | *Actinobacteria* | 2905150 | 75.39 | 31 | 2745 | 100 |
| LAC_ARCH01 | anamox2_Methanosarcina_thermophila_41_9_curated | *Euryarchaeota* | 2959285 | 41.19 | 82 | 2703 | 35.55 |
| LAC_ARM01 | LAC_NA06_Fimbriimonas_ginsengisoli_61_14_curated | *Armatimonadetes* | 2673573 | 61.04 | 54 | 2542 | 88.45 |
| LAC_BAC01 | anamox1_Bacteria_33_9_curated | *Microgenomates* | 775494 | 32.38 | 45 | 793 | 70.85 |
| LAC_BAC02 | anamox1_Bacteria_45_8_curated | *Ignavibacteria* | 2579669 | 45 | 166 | 2221 | 87.77 |
| LAC_BAC03 | anamox1_Bacteria_50_18_curated | *TM7* | 820566 | 49.55 | 22 | 874 | 77.9 |
| LAC_BAC04 | anamox1_Bacteria_53_17_curated | *Deltaproteobacteria* | 3465172 | 53.24 | 12 | 2895 | 100 |
| LAC_BAC05 | anamox1_Bacteria_55_18_curated | Bacteria | 4100779 | 54.56 | 216 | 3365 | 98.28 |
| LAC_BAC06 | anamox1_Bacteria_57_32_curated | *Lentisphaerae* | 2939326 | 56.91 | 76 | 2571 | 95.69 |
| LAC_BAC07 | anamox1_Bacteria_57_5_curated | *Chloroflexi* | 3332363 | 56.3 | 813 | 3846 | 68.62 |
| LAC_BAC08 | anamox1_Bacteria_57_9_curated | *Planctomycetes* | 3469651 | 57.28 | 74 | 3012 | 93.65 |
| LAC_BAC09 | anamox1_Bacteria_64_7_curated | *Planctomycetes* | 3988790 | 63.42 | 306 | 3679 | 93.03 |
| LAC_BAC10 | anamox1_Bacteria_65_5_curated | *Planctomycetes* | 2089721 | 64.54 | 434 | 2237 | 51.28 |
| LAC_BAC11 | anamox1_Bacteria_72_15_curated | *Acidobacteria* | 2937143 | 72.45 | 24 | 2507 | 87.93 |
| LAC_BAC12 | anamox2_Bacteria_34_5_curated | *Microgenomates* | 434538 | 33.34 | 88 | 508 | 50.78 |
| LAC_BAC13 | anamox2_Bacteria_61_6_curated | *Armatimonadetes* | 2415824 | 60.87 | 385 | 2646 | 65.91 |
| LAC_BAC14 | anamox2_Bacteria_68_6_curated | Bacteria | 2291336 | 67.96 | 460 | 2621 | 64.03 |
| LAC_BAC15 | anamox3_Bacteria_50_5_curated | *Ignavibacteria* | 1042006 | 49.54 | 203 | 1076 | 42.87 |
| LAC_BAC16 | anamox3_Bacteria_66_7_curated | *Verrucomicrobia* | 2266342 | 65.35 | 286 | 2150 | 81.9 |
| LAC_BAC17 | anamox3_Bacteria_67_13_curated | Bacteria | 1642722 | 66.5 | 47 | 1442 | 67.24 |
| LAC_BAC18 | anamox3_Bacteria_67_15_curated | *Chloroflexi* | 2765727 | 66.48 | 144 | 2980 | 95.69 |
| LAC_BAC19 | anamox4_Bacteria_63_7_curated | *Chloroflexi* | 3451341 | 62.64 | 351 | 3276 | 93.1 |
| LAC_BAC20 | anamox4_Bacteria_69_43_curated | *Deltaproteobacteria* | 3827900 | 69.37 | 15 | 3388 | 86.21 |
| LAC_BAC21 | LAC_NA06_Bacteria_60_18_curated | *Deltaproteobacteria* | 3689918 | 59.98 | 50 | 3221 | 94.83 |
| LAC_BAC22 | LAC_NA07_Bacteria_38_171_curated | *Ignavibacteria* | 2383463 | 37.56 | 22 | 2157 | 98.28 |
| LAC_BAC23 | LAC_NA07_Bacteria_70_305_curated | *Planctomycetes* | 2792777 | 70.13 | 54 | 2465 | 98.28 |

| | | | | | | | |
|---|---|---|---|---|---|---|---|
| LAC_BAC24 | LAC_NA07_Bacteria_71_12_curated | *Chloroflexi* | 2770642 | 70.73 | 206 | 2877 | 81.41 |
| LAC_BACT01 | anamox1_Bacteroidetes_38_5_curated | *Bacteroidetes* | 1414251 | 37.34 | 293 | 1501 | 62.96 |
| LAC_BACT02 | anamox1_Bacteroidetes_39_16_curated | *Bacteroidetes* | 2896628 | 38.84 | 40 | 2472 | 100 |
| LAC_BACT03 | anamox1_Bacteroidetes_63_11_curated | *Bacteroidetes* | 3356307 | 63.21 | 73 | 2915 | 100 |
| LAC_BACT04 | anamox1_Sphingobacteriales_42_27_curated | *Bacteroidetes* | 3490519 | 42 | 95 | 3068 | 100 |
| LAC_BACT05 | anamox1_Sphingobacteriales_43_8_curated | *Bacteroidetes* | 3156182 | 42.95 | 130 | 2826 | 97.41 |
| LAC_BACT06 | anamox2_BJP_IG2103_Bacteroidetes_37_22_46_7_curated | *Bacteroidetes* | 2490972 | 45.52 | 190 | 2226 | 93.1 |
| LAC_BACT07 | anamox2_Sphingobacteriales_41_11_curated | *Bacteroidetes* | 2768577 | 41.16 | 129 | 2490 | 87.07 |
| LAC_BACT08 | anamox3_Bacteroidetes_39_15_curated | *Bacteroidetes* | 2533542 | 39.26 | 23 | 2188 | 100 |
| LAC_BACT09 | anamox3_Burkholderiales_71_6_curated | *Bacteroidetes* | 1367674 | 69.86 | 648 | 1904 | 32.85 |
| LAC_BACT10 | anamox3_Sphingobacteriales_44_6_curated | *Bacteroidetes* | 3033899 | 42.77 | 512 | 2768 | 73.9 |
| LAC_BACT11 | anamox3_Sphingobacteriales_50_9_curated | *Bacteroidetes* | 4305233 | 49.73 | 216 | 2999 | 99.14 |
| LAC_BACT12 | anamox4_Bacteroidetes_40_74_curated | *Bacteroidetes* | 2627547 | 39.91 | 15 | 2219 | 100 |
| LAC_BACT13 | LAC_NA06_Bacteroidetes_30_9_curated | *Bacteroidetes* | 2233050 | 29.92 | 225 | 2055 | 85.06 |
| LAC_CHLX01 | anamox1_Bacteria_56_37_curated | *Chloroflexi* | 4970250 | 55.88 | 43 | 4404 | 94.83 |
| LAC_CHLX02 | anamox1_Chloroflexi_52_59_curated | *Chloroflexi* | 2864537 | 52.47 | 96 | 2759 | 100 |
| LAC_CHLX03 | anamox2_Chloroflexi_60_8_curated | *Chloroflexi* | 2023353 | 60.06 | 111 | 1885 | 47.49 |
| LAC_CHLX04 | anamox3_Chloroflexi_59_6_curated | *Chloroflexi* | 1407767 | 53.68 | 351 | 1507 | 40.22 |
| LAC_CHLX05 | anamox3_Chloroflexi_68_6_curated | *Chloroflexi* | 2294191 | 67.35 | 479 | 2330 | 53.92 |
| LAC_CHLX06 | anamox4_Chloroflexi_66_15_curated | *Chloroflexi* | 4383590 | 65.96 | 133 | 3602 | 91.22 |
| LAC_CHLX07 | LAC_NA06_Anaerolineales_42_27_curated | *Chloroflexi* | 2409642 | 41.78 | 210 | 2380 | 93.1 |
| LAC_CHLX08 | LAC_NA06_Chloroflexi_57_14_curated | *Chloroflexi* | 2719486 | 57.25 | 159 | 2567 | 86.91 |
| LAC_CHLX09 | LAC_NA06_sub_Chloroflexi_59_14_curated | *Chloroflexi* | 2594752 | 59.45 | 164 | 2359 | 91.22 |
| LAC_CHLX10 | LAC_NA06_sub_Chloroflexi_61_22_curated | *Chloroflexi* | 2755473 | 61.22 | 170 | 2707 | 96.55 |
| LAC_CHLX11 | LAC_NA07_Caldilinea_aerophila_61_12_curated | *Chloroflexi* | 3885866 | 60.59 | 83 | 3168 | 80.88 |
| LAC_CHLX12 | LAC_NA07_Chloroflexi_57_23_curated | *Chloroflexi* | 6358497 | 57.22 | 93 | 5201 | 89.66 |
| LAC_CHLX13 | LAC_NA07_Chloroflexi_57_9_curated | *Chloroflexi* | 2081669 | 55.57 | 325 | 2239 | 71.76 |
| LAC_CHLX14 | LAC_NA07_Chloroflexi_58_12_curated | *Chloroflexi* | 2711149 | 57.43 | 154 | 2459 | 60.42 |
| LAC_CHLX15 | LAC_NA07_Chloroflexi_60_59_curated | *Chloroflexi* | 3858643 | 60.1 | 13 | 3478 | 94.83 |
| LAC_CHLX16 | LAC_NA07_Chloroflexi_65_58_curated | *Chloroflexi* | 3316034 | 65.08 | 327 | 3017 | 87.93 |
| LAC_CHLX17 | LAC_NA07_Chloroflexi_67_63_curated | *Chloroflexi* | 3446202 | 66.61 | 65 | 2852 | 96.55 |
| LAC_CHLX18 | LAC_NA07_RBG_16_RIF_CHLX_72_14_curated_75_20_curated | *Chloroflexi* | 2506614 | 74.69 | 63 | 2330 | 91.38 |
| LAC_CLO01 | anamox1_Candidatus_Cloacimonas_acidaminovorans_38_6_curated | Bacteria | 1192038 | 35.28 | 246 | 1132 | 56.47 |
| LAC_D-T01 | anamox1_Truepera_radiovictrix_71_5_curated | *Deinococcus Thermus* | 435656 | 69.12 | 317 | 694 | 24.49 |
| LAC_D-T02 | LAC_NA07_Truepera_radiovictrix_72_29_curated | *Deinococcus Thermus* | 1353035 | 72.21 | 165 | 1373 | 82.6 |
| LAC_DADA01 | LAC_NA06_RIFCSPHIGHO2_12_FULL_Dadabacteria_53_21_curated_58_6_curated | *Dadabacteria* | 869333 | 55.01 | 539 | 1358 | 33.68 |
| LAC_GMT01 | anamox1_Gemmatimonas_aurantiaca_57_6_curated | *Gemmatimonadetes* | 2631895 | 56.94 | 255 | 2549 | 91.22 |
| LAC_IGN01 | anamox1_RBG_16_Ignavibacteria_36_9_curated_35_5_curated | *Ignavibacteria* | 1163158 | 34.03 | 347 | 1374 | 70.14 |
| LAC_IGN02 | anamox2_Ignavibacteriales_33_72_curated | *Ignavibacteria* | 3331053 | 33.19 | 161 | 2995 | 98.28 |

| | | | | | | | |
|---|---|---|---|---|---|---|---|
| LAC_IGN03 | anamox2_Ignavibacteriales_33_9_curated | *Ignavibacteria* | 2936140 | 33.09 | 166 | 2753 | 86.05 |
| LAC_IGN04 | anamox2_Ignavibacteriales_41_12_curated | *Ignavibacteria* | 3156563 | 41.32 | 42 | 2805 | 96.55 |
| LAC_IGN05 | anamox2_sub_Ignavibacterium_album_33_16_curated | *Ignavibacteria* | 3391206 | 33.43 | 195 | 3029 | 97.81 |
| LAC_IGN06 | anamox3_BJP_IG2069_Ignavibacteriae_38_11_30_7_curated | *Ignavibacteria* | 1803433 | 30.38 | 132 | 1600 | 73.51 |
| LAC_IGN07 | anamox3_sub_Ignavibacteriales_42_14_curated | *Ignavibacteria* | 3163923 | 42.21 | 22 | 2498 | 96.55 |
| LAC_MIC01 | anamox2_Microgenomates_45_6_curated | *Microgenomates* | 740215 | 44.28 | 99 | 904 | 73.35 |
| LAC_MIC02 | anamox2_Microgenomates_49_6_curated | *Microgenomates* | 950204 | 49.23 | 109 | 1082 | 68.42 |
| LAC_MIC03 | anamox2_Roizmanbacteria_38_11_curated | *Microgenomates* | 663231 | 37.62 | 14 | 693 | 58.31 |
| LAC_MIC04 | anamox4_Microgenomates_48_8_curated | *Microgenomates* | 999881 | 47.64 | 49 | 1055 | 80.02 |
| LAC_MIC05 | LAC_NA06_Microgenomates_41_17_curated | *Microgenomates* | 1110978 | 41.17 | 7 | 1216 | 80.88 |
| LAC_MIC06 | LAC_NA07_Roizmannbacteria_52_60_curated | *Microgenomates* | 957307 | 52.33 | 6 | 1013 | 73.98 |
| LAC_NIT01 | anamox4_Candidatus_Nitrospira_defluvii_60_9_curated | *Microgenomates* | 3085099 | 60.3 | 94 | 3084 | 95.69 |
| LAC_OMN01 | anamox3_Omnitrophica_63_14_curated | *Omnitrophica* | 365510 | 62.81 | 4 | 402 | 25.86 |
| LAC_PLT01 | anamox1_Planctomycetia_64_8_curated | *Planctomycetes* | 4138393 | 63.58 | 783 | 3420 | 87.62 |
| LAC_PLT02 | anamox4_sub_Candidatus_Brocadia_sinica_42_75_curated | *Planctomycetes* | 3107335 | 42.29 | 64 | 2859 | 98.28 |
| LAC_PROT01 | anamox1_Betaproteobacteria_69_7_curated | *Betaproteobacteria* | 2106569 | 66.39 | 531 | 2636 | 66.55 |
| LAC_PROT02 | anamox1_Burkholderiales_70_40_curated | *Betaproteobacteria* | 3601855 | 70.09 | 60 | 3356 | 100 |
| LAC_PROT03 | anamox1_Burkholderiales_71_17_curated | *Betaproteobacteria* | 3639459 | 70.81 | 114 | 3448 | 99.22 |
| LAC_PROT04 | anamox1_Gammaproteobacteria_64_6_curated | *Gammaproteobacteria* | 2447777 | 64.02 | 391 | 2594 | 84.25 |
| LAC_PROT05 | anamox1_Lysobacter_67_10_curated | *Gammaproteobacteria* | 783854 | 66.76 | 125 | 840 | 34.64 |
| LAC_PROT06 | anamox1_Nitrosomonas_europaea_50_14_curated | *Betaproteobacteria* | 2128781 | 50.43 | 37 | 2006 | 98.28 |
| LAC_PROT07 | anamox1_Proteobacteria_65_15_curated | *Gammaproteobacteria* | 2240508 | 65.45 | 348 | 2476 | 58.59 |
| LAC_PROT08 | anamox1_Proteobacteria_67_8_curated | *Gammaproteobacteria* | 2912885 | 66.93 | 203 | 2910 | 81.27 |
| LAC_PROT09 | anamox1_Rhodocyclales_69_13_curated | *Betaproteobacteria* | 2263347 | 68.53 | 185 | 2364 | 96.55 |
| LAC_PROT10 | anamox2_Burkholderiales_67_5_curated | *Betaproteobacteria* | 1246117 | 66.28 | 298 | 1434 | 38.05 |
| LAC_PROT11 | anamox2_Burkholderiales_68_9_curated | *Betaproteobacteria* | 3527974 | 68.25 | 181 | 3452 | 91.22 |
| LAC_PROT12 | anamox2_Burkholderiales_70_13_curated | *Betaproteobacteria* | 3074974 | 70.33 | 39 | 2841 | 84.48 |
| LAC_PROT13 | anamox2_Gammaproteobacteria_67_23_curated | *Gammaproteobacteria* | 2663638 | 67.06 | 116 | 2584 | 79.31 |
| LAC_PROT14 | anamox2_Hydrogenophilales_66_19_curated | *Betaproteobacteria* | 2257110 | 65.94 | 122 | 2331 | 91.95 |
| LAC_PROT15 | anamox2_Myxococcales_71_5_curated | *Deltaproteobacteria* | 1434218 | 70.21 | 495 | 1687 | 30.21 |
| LAC_PROT16 | anamox2_Rhizobiales_67_45_curated | *Alphaproteobacteria* | 4837226 | 66.56 | 24 | 4668 | 100 |
| LAC_PROT17 | anamox2_Xanthomonadales_68_9_curated | *Gammaproteobacteria* | 1242229 | 67.63 | 21 | 1168 | 50 |
| LAC_PROT18 | anamox2_Xanthomonadales_70_8_curated | *Gammaproteobacteria* | 2605543 | 68.54 | 346 | 2555 | 82.76 |
| LAC_PROT19 | anamox3_Nitrosomonas_europaea_51_28_curated | *Betaproteobacteria* | 2372894 | 50.54 | 93 | 2302 | 98.12 |
| LAC_PROT20 | anamox3_Proteobacteria_68_11_curated | *Gammaproteobacteria* | 3218375 | 68.07 | 66 | 3078 | 100 |
| LAC_PROT21 | anamox4_Alphaproteobacteria_58_14_curated | *Alphaproteobacteria* | 2593178 | 57.86 | 5 | 2558 | 98.28 |
| LAC_PROT22 | anamox4_Gammaproteobacteria_67_14_curated | *Gammaproteobacteria* | 3265689 | 66.91 | 73 | 3049 | 98.28 |
| LAC_PROT23 | anamox4_Nitrosomonas_eutropha_48_11_curated | *Betaproteobacteria* | 1880399 | 48.46 | 103 | 1862 | 94.83 |
| LAC_PROT24 | LAC_NA06_Betaproteobacteria_71_7_curated | *Betaproteobacteria* | 4060980 | 69.45 | 1841 | 5695 | 69.75 |

| | | | | | | | |
|---|---|---|---|---|---|---|---|
| LAC_PROT25 | LAC_NA06_Burkholderiales_73_13_curated | *Betaproteobacteria* | 3708868 | 72.67 | 335 | 3469 | 94.51 |
| LAC_PROT26 | LAC_NA06_Rhizobiales_66_24_curated | *Alphaproteobacteria* | 3228463 | 66.32 | 148 | 3270 | 98.28 |
| LAC_PROT27 | LAC_NA07_Burkholderiales_70_312_curated | *Betaproteobacteria* | 2688511 | 69.75 | 290 | 2827 | 93.32 |
| LAC_PROT28 | LAC_NA07_Proteobacteria_68_32_curated | *Gammaproteobacteria* | 2747459 | 68.4 | 14 | 2541 | 98.28 |
| LAC_PROT29 | LAC_NA07_Rhodobacterales_68_7_curated | *Alphaproteobacteria* | 1308110 | 67.47 | 473 | 1651 | 52.93 |
| LAC_PROT30 | LAC_NA07_Rhodocyclales_67_14_curated | *Betaproteobacteria* | 2723825 | 66.65 | 212 | 2909 | 92.95 |
| LAC_SCH01 | LAC_NA06_Candidatus_Saccharibacteria_40_6_curated | *TM7* | 463567 | 39.93 | 148 | 580 | 58.46 |
| LAC_SPR01 | anamox1_Turneriella_parva_44_7_curated | *Spirochaetes* | 2667359 | 44.12 | 240 | 2748 | 92.24 |
| LAC_VER01 | anamox2_Verrucomicrobia_58_8_curated | *Verrucomicrobia* | 3592680 | 57.68 | 250 | 3293 | 87.93 |
| LAC_VER02 | anamox2_Verrucomicrobia_62_8_curated | *Verrucomicrobia* | 3685167 | 61.78 | 161 | 3134 | 98.28 |
| LAC_VER03 | anamox3_Opitutus_terrae_67_4_curated | *Verrucomicrobia* | 780321 | 65.72 | 503 | 1046 | 24.42 |
| LAC_VER04 | anamox3_Pedosphaera_parvula_66_5_curated | *Verrucomicrobia* | 1233438 | 65.98 | 543 | 1384 | 56 |
| LAC_VER05 | anamox3_Verrucomicrobia_59_12_curated | *Verrucomicrobia* | 2712092 | 59.27 | 50 | 2592 | 96.55 |

**Temporal strain shifts in the bioreactor.** Between Days 82 and 328, we binned 2-3 partial bins of the *Brocadia* sp (at each time point). These partial bins were separated by their coverage peaks, indicating strain variation. The presence of multiple strains at some of the time points was verified by the dereplication process. The *Brocadia* sp. cluster contained complete genome bins of the two strains: anamox1_Candidatus_Brocadia_sinica_42_73_curated (strain 1) and LAC_PLT02 (strain 2 and the representative genome of the anammox bacterium). The first genome was binned from the source community and can be found up to Day 166 in the reactor. The second genome is represented from Day 284 and onwards. This strain is the only *Brocadia* sp. bacterium (and anammox bacterium) present at stable state in Day 437.

Apart from *Brocadia* sp., we observed strain shift in eight other bacteria around the same time. Several are noteworthy: LAC_IGN05, which shifted with anamox4_Ignavibacterium_album_34_177_curated, was the most dominant bacteria during the destabilization period. Apart from this bacterium, three other bacteria were also associated with the destabilization. It seems that the biomass amendment introduced not only an anammox bacterium that came to dominate the community, but also strong destabilizing bacteria.

anamox2_sub_Burkholderiales_70_25_curated shifted into LAC_PROT27, and LAC_MIC02 shifted to LAC_NA07_Microgenomates_50_23_curated. The first bacterium was one of three bacteria consistently in the top ten abundant bacteria. Its abundance was highest when the anammox bioreactor was performing well. The second bacterium exhibited a three-fold increase in abundance between Days 82 and 437. These two bacteria were left outliers (on the x axis) of Group D in the nMDS (Figure 4B). While the pairwise correlation between the two bacteria is not the highest among Group D, the results of the nMDS and strain shift might suggest that the two are a host-CPR pair.

**Table S2** Genome parameters for the strain bacteria from the anammox reactor

| **Genome** | **Taxonomy (ggKbase)** | **length (bp)** | **GC%** | **contigs** | **features** | **Completeness** |
|---|---|---|---|---|---|---|
| anamox1_Candidatus_Brocadia_sinica_42_73_curated | *Candidatus Brocadia sinica, Candidatus Brocadia, Candidatus Brocadiales, Planctomycetia, Planctomycetes, Bacteria* | 3182191 | 42.3 | 140 | 2861 | 98.28 |
| anamox2_Chloroflexi_65_7_curated | *Chloroflexi, Bacteria* | 2846499 | 64 | 427 | 2818 | 68.5 |
| anamox2_sub_Burkholderiales_70_25 _curated | *Burkholderiales, Betaproteobacteria, Proteobacteria, Bacteria* | 1637210 | 69.92 | 260 | 1745 | 86.21 |
| anamox3_Actinobacteria_74_6_curated | *Actinobacteria, Actinobacteria, Bacteria* | 1306370 | 70.77 | 293 | 1474 | 33.9 |
| anamox3_Truepera_radiovictrix_72_6 _curated | *Truepera radiovictrix, Truepera, Deinococcales, Deinococci, Deinococcus-Thermus, Bacteria* | 1087782 | 69.47 | 428 | 1343 | 47.82 |
| anamox4_Betaproteobacteria_69_7 _curated | *Betaproteobacteria, Proteobacteria, Bacteria* | 2004013 | 69.34 | 569 | 2383 | 59.72 |
| anamox4_Ignavibacterium_album_34_177_curated | *Ignavibacterium album, Ignavibacterium, Ignavibacteriales, Ignavibacteria, Ignavibacteriae, Bacteria* | 1414754 | 33.55 | 464 | 1618 | 63.32 |
| LAC_NA06_Rubrivivax_70_13_curated | *Rubrivivax, Burkholderiales, Betaproteobacteria, Proteobacteria, Bacteria* | 1206889 | 70.2 | 103 | 1213 | 70.61 |
| LAC_NA07_Microgenomates_50_23 _curated | *Microgenomates, Bacteria* | 947146 | 49.7 | 96 | 1047 | 66.74 |

**Table S3** assignment of genomes to different clusters

| Genome | Abundance Group | Metabolic Group | Association grouping |
|---|---|---|---|
| LAC_ACD01 | A | NA | NA |
| LAC_ACD02 | A | NA | NA |
| LAC_ACD03 | C | ɣ | AA |
| LAC_ACD04 | A | NA | NA |
| LAC_ACD05 | B | ɣ | AA |
| LAC_ACD06 | D | ɣ | AA |
| LAC_ACD07 | D | NA | NA |
| LAC_ACT01 | C | NA | NA |
| LAC_ACT02 | D | α | AA |
| LAC_ACT03 | B | NA | NA |
| LAC_ACT04 | D | α | AA |
| LAC_ARCH01 | C | NA | NA |
| LAC_ARM01 | D | ɣ | AA |
| LAC_BAC01 | A | β | SA |
| LAC_BAC02 | A | δ | SA |
| LAC_BAC03 | A | β | SA |
| LAC_BAC04 | A | ɣ | SA |
| LAC_BAC05 | A | ɣ | SA |
| LAC_BAC06 | A | ɣ | SA |
| LAC_BAC07 | A | NA | NA |
| LAC_BAC08 | A | ɣ | SA |
| LAC_BAC09 | A | ɣ | SA |
| LAC_BAC10 | A | NA | NA |
| LAC_BAC11 | A | ɣ | SA |
| LAC_BAC12 | C | NA | NA |
| LAC_BAC13 | C | NA | NA |
| LAC_BAC14 | C | NA | NA |
| LAC_BAC15 | A | NA | NA |
| LAC_BAC16 | A | ɣ | SA |
| LAC_BAC17 | A | NA | NA |
| LAC_BAC18 | A | α | SA |
| LAC_BAC19 | B | α | AA |
| LAC_BAC20 | B | ɣ | AA |
| LAC_BAC21 | A | ɣ | AA |
| LAC_BAC22 | D | δ | AA |
| LAC_BAC23 | D | ɣ | AA |

| | | | |
|---|---|---|---|
| LAC_BAC24 | B | α | AA |
| LAC_BACT01 | A | NA | NA |
| LAC_BACT02 | A | δ | SA |
| LAC_BACT03 | C | δ | SA |
| LAC_BACT04 | B | ɣ | AA |
| LAC_BACT05 | A | ɣ | SA |
| LAC_BACT06 | C | δ | SA |
| LAC_BACT07 | C | ɣ | SA |
| LAC_BACT08 | A | δ | SA |
| LAC_BACT09 | A | NA | NA |
| LAC_BACT10 | A | ɣ | SA |
| LAC_BACT11 | A | δ | SA |
| LAC_BACT12 | A | δ | AA |
| LAC_BACT13 | B | δ | SA |
| LAC_CHLX01 | B | α | AA |
| LAC_CHLX02 | A | α | SA |
| LAC_CHLX03 | C | NA | NA |
| LAC_CHLX04 | B | NA | NA |
| LAC_CHLX05 | A | NA | NA |
| LAC_CHLX06 | B | α | AA |
| LAC_CHLX07 | B | α | AA |
| LAC_CHLX08 | D | α | AA |
| LAC_CHLX09 | B | α | AA |
| LAC_CHLX10 | D | α | AA |
| LAC_CHLX11 | D | α | AA |
| LAC_CHLX12 | D | α | AA |
| LAC_CHLX13 | D | α | AA |
| LAC_CHLX14 | D | NA | NA |
| LAC_CHLX15 | D | α | AA |
| LAC_CHLX16 | D | α | AA |
| LAC_CHLX17 | D | α | AA |
| LAC_CHLX18 | D | α | AA |
| LAC_CLO01 | A | NA | NA |
| LAC_D-T01 | B | NA | NA |
| LAC_D-T02 | B | α | AA |
| LAC_DADA01 | B | NA | NA |
| LAC_GMT01 | A | ɣ | SA |
| LAC_IGN01 | A | δ | SA |
| LAC_IGN02 | C | δ | AA |

| | | | |
|---|---|---|---|
| LAC_IGN03 | C | δ | SA |
| LAC_IGN04 | C | ɣ | SA |
| LAC_IGN05 | B | δ | AA |
| LAC_IGN06 | B | δ | SA |
| LAC_IGN07 | A | ɣ | AA |
| LAC_MIC01 | D | β | AA |
| LAC_MIC02 | D | NA | NA |
| LAC_MIC03 | C | NA | NA |
| LAC_MIC04 | B | β | AA |
| LAC_MIC05 | B | β | AA |
| LAC_MIC06 | D | β | AA |
| LAC_NIT01 | A | ɣ | SA |
| LAC_OMN01 | A | NA | NA |
| LAC_PLT01 | A | ɣ | SA |
| LAC_PLT02 | D | ɣ | AA |
| LAC_PROT01 | B | NA | NA |
| LAC_PROT02 | A | ε | AA |
| LAC_PROT03 | A | ε | AA |
| LAC_PROT04 | A | ɣ | SA |
| LAC_PROT05 | A | NA | NA |
| LAC_PROT06 | A | ε | SA |
| LAC_PROT07 | A | NA | NA |
| LAC_PROT08 | A | ε | SA |
| LAC_PROT09 | A | ε | SA |
| LAC_PROT10 | C | NA | NA |
| LAC_PROT11 | C | ε | SA |
| LAC_PROT12 | C | ε | SA |
| LAC_PROT13 | C | ε | SA |
| LAC_PROT14 | C | ε | SA |
| LAC_PROT15 | C | NA | NA |
| LAC_PROT16 | C | ε | SA |
| LAC_PROT17 | C | NA | NA |
| LAC_PROT18 | B | ε | AA |
| LAC_PROT19 | A | ε | SA |
| LAC_PROT20 | A | ε | SA |
| LAC_PROT21 | D | ε | AA |
| LAC_PROT22 | B | ε | AA |
| LAC_PROT23 | A | ε | AA |
| LAC_PROT24 | B | NA | NA |

| | | | |
|---|---|---|---|
| LAC_PROT25 | B | ε | SA |
| LAC_PROT26 | D | ε | AA |
| LAC_PROT27 | D | ε | AA |
| LAC_PROT28 | D | ε | AA |
| LAC_PROT29 | D | NA | NA |
| LAC_PROT30 | C | ε | AA |
| LAC_SCH01 | B | NA | NA |
| LAC_SPR01 | A | ɣ | SA |
| LAC_VER01 | C | ɣ | SA |
| LAC_VER02 | A | ɣ | SA |
| LAC_VER03 | A | NA | NA |
| LAC_VER04 | A | NA | NA |
| LAC_VER05 | A | ɣ | SA |

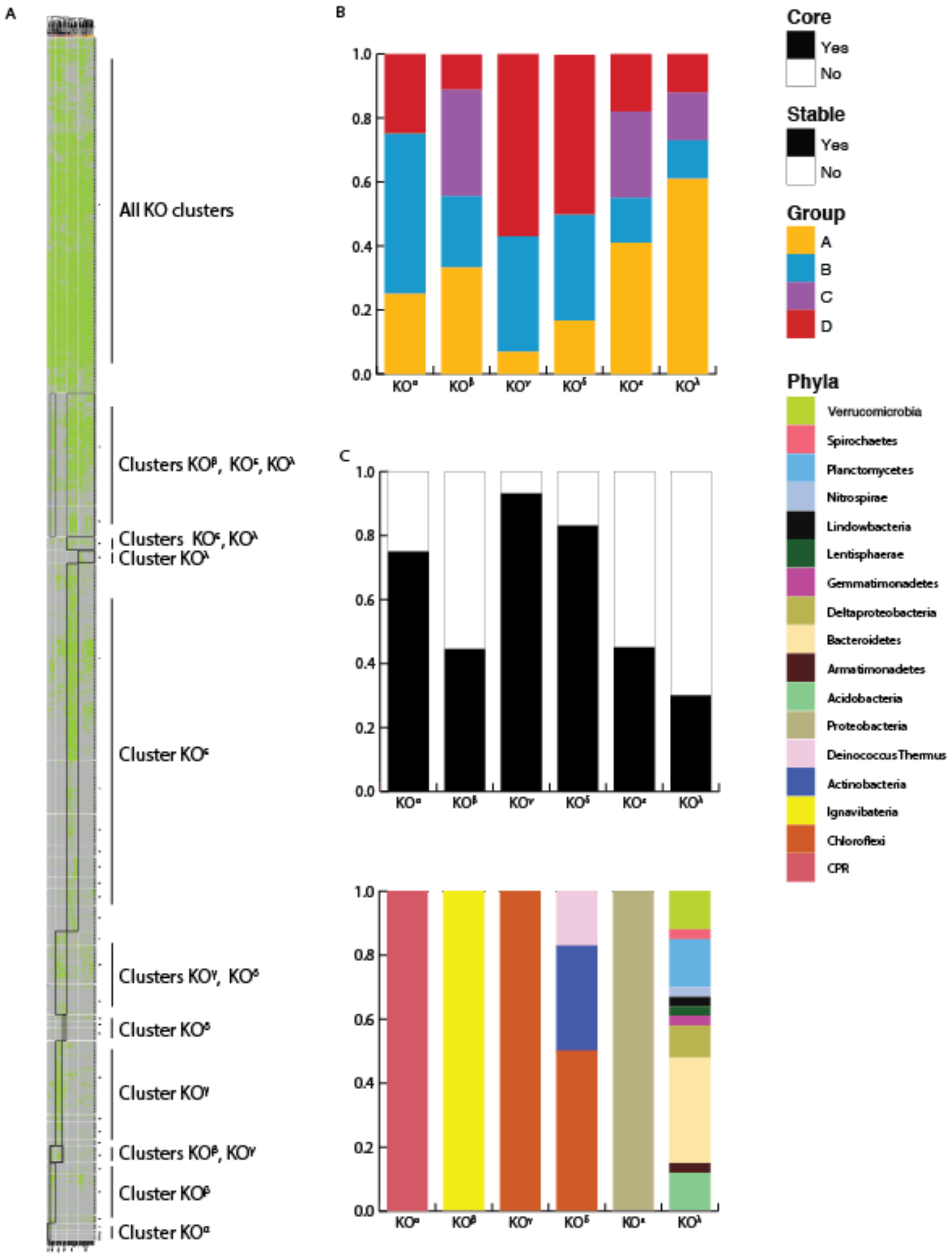

**Figure S1 Analysis of community metabolism by gene presence.** Genes were annotated against the KEGG database. A) A presence/absence-based clustering of both genomes and KOs using Jaccard distance and complete clustering method. The dual clustering forms "blocks" across rows and columns, and blocks that are representative of a genome cluster are presented. Black squares mark the representative blocks and on the right, the relevant cluster is mentioned. B) Fractionation of the KO cluster by abundance grouping (A-D). C) Fractionation of KO clusters by SA and AA groups. Both panels C and D indicate that clusters KOγ and KOδ are highly associated with stable state, while cluster KO$\lambda$ is the most associated with the sludge.  D) Fractionation of KO clusters by taxonomy. Most KO clusters have a strong phylogenetic signal, with four of six having members of a single phylum.

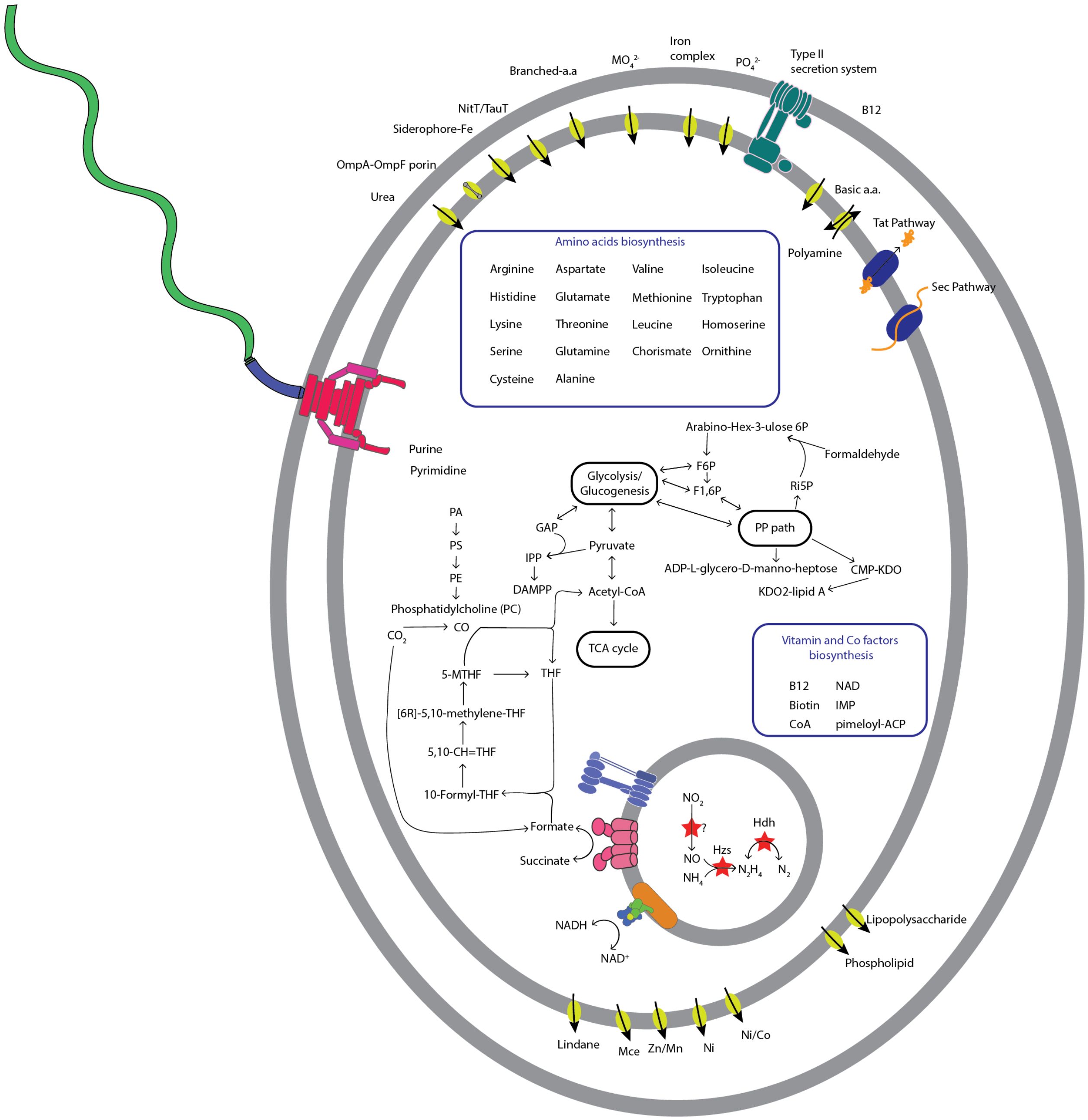

**Figure S2 Reduced metabolic map of the *Brocadia* sp. Bacterium.** The bacterium has the enzymes needed for the anammox process, and fixes carbon via the Wood-Ljungdahl pathway. It can synthesize 18/20 essential aa., and is the only bacterium in the reactor that can synthesize vitamin B12.

**Electron transfer pathways in the community**. Apart from nitrogen reduction, another common anaerobic respiration pathway within the community is acetate fermentation (genes detected in 60% of the genomes). This process was more common in AA (69%) bacteria than in SA (51%) bacteria. Ni-Fe Hydrogenase was present in 31% of the genomes, but was most common among the *Chloroflexi* of group α (87% and 48% of all occurrences of hydrogenases found) (Figure 6B).

The majority of bacteria in the bioreactor are potentially facultative aerobes (58%). All had high affinity complex IV, which differed between AA and SA bacteria. In the AA bacteria, the bd type was found in all aerobic members of group α (one also has a cbb3 type) and the *Ignavibacteria*, and the cbb3 type occurred mostly in *Proteobacteria*. For the SA bacteria, the cbb3 type was found in 24/25 aerobes and the bd-type was only found in 6/25 (only in one bacterium it is the sole variant). Complex III, which is also essential to aerobic respiration, was only found in 14 *Proteobacteria*, one *Actinobacterium*, and one *Chloroflexi*. It is possible that other bacteria have an alternative Complex III [30] that cannot be found by current KEGG annotations. Complexes I/II were found in nearly all of the bacteria, except CPR. Only five bacteria lacked the F-type ATPase; two had the V-type ATPase instead.

**Central carbon metabolism in the community**. It is likely that nearly all bacteria (98%) in the bioreactor can oxidize sugar by glycolysis (Figure 6A and E). Fewer bacteria (69%) have the pentose phosphate pathway (PPP). Acetyl-CoA could be synthesized from pyruvate (90% general, 98% AA, and 81% SA), or by beta-oxidation (49% general, 57% AA, and 43% SA). The majority of bacteria have the full TCA cycle (84%, or 88% after excluding CPR). A possible major carbon source for the bacteria in the bioreactor are amino acids (aa.), with 95% being able to incorporate aa. into their central carbon metabolism. The most common aa. (aspartate) could be converted into oxaloacetate and fed into the TCA cycle. Three aa. (serine, alanine, and cysteine) could be converted to pyruvate. Of these, only cysteine conversion is unidirectional, so aa., as a carbon source, cannot be ascertained. Group α had additional genes that support a reliance on proteins for their metabolism (Figure 6B). They also had a set of peptidases, as well as multiple transporters covering all forms of aa., peptides, and polyamines.

Some metabolic groups could use aa. as precursors for synthesis of other metabolites. Glutamate and histidine could be converted to PRPP, and glutamine to pyrimidines (Figure 6A). Groups ɣ and ε could use aspartate to synthesize $NAD^+$, and glutamine to synthesis IMP (Figure 6C). $NAD^+$ and IMP could not be synthesized by all of the bacteria, indicating that there are potential metabolic interdependencies in the community. Members of group ε (Figure 6D) could use leucine as a precursor to acetyl-CoA, lysine for acetoacetyl-CoA, glutamate for glutathione, and chorismate for ubiquinone. The last two could only be synthesized by group ε, indicating additional potential metabolic interdependencies in the community.

**Table S4** Diversity indices of the anammox community

A) Alpha diversity (Shannon) for pairwise timepoints

| | Day 0 | Day 82 | Day 166 | Day 284 | Day 328 | Day 437 |
|---|---|---|---|---|---|---|
| **Day 0** | 29.23 | 22.81 | 27.89 | 14.28 | 18.59 | 18.59 |
| **Day 82** | 22.81 | 17.81 | 21.77 | 11.15 | 14.52 | 14.52 |
| **Day 166** | 27.89 | 21.77 | 26.62 | 13.63 | 17.75 | 17.75 |
| **Day 284** | 14.28 | 11.15 | 13.63 | 6.98 | 9.09 | 9.09 |
| **Day 328** | 24.17 | 18.87 | 23.07 | 11.81 | 15.38 | 15.38 |
| **Day 437** | 18.59 | 14.52 | 17.75 | 9.09 | 11.83 | 11.83 |

B) Beta diversity for pairwise timepoints

| | DAY 0 | DAY 82 | DAY 166 | DAY 284 | DAY 328 | DAY 437 |
|---|---|---|---|---|---|---|
| **DAY 0** | 1.00 | 1.41 | 1.08 | 1.39 | 1.55 | 1.55 |
| **DAY 82** | 1.41 | 1.00 | 1.21 | 1.14 | 1.18 | 1.18 |
| **DAY 166** | 1.08 | 1.21 | 1.00 | 1.17 | 1.31 | 1.31 |
| **DAY 284** | 1.39 | 1.14 | 1.17 | 1.00 | 1.12 | 1.12 |
| **DAY 328** | 1.37 | 1.17 | 1.20 | 1.14 | 1.11 | 1.11 |
| **DAY 437** | 1.55 | 1.18 | 1.31 | 1.12 | 1.00 | 1.00 |

**Table S5** Synthesis and transport of metabolites in the MO groups. *Brocadia sp.* is considered separately from its MO grouping. Green denotes >50% of the group have the ability to synthesis/transport. Yellow denotes >40% of the group have the ability to synthesis/transport. Red denotes <40% of the group have the ability to synthesis/transport.

A) Synthesis of aa.

| Module | Amino acid | MOα | MOγ | Brocadia | MOβ | MOδ | MOε |
|---|---|---|---|---|---|---|---|
| By KO | arginine | Green | Green | Green | Red | Red | Green |
| M00026 | histidine | Green | Green | Green | Red | Red | Green |
| M00016/M00526 | lysine | Green | Green | Green | Red | Red | Green |
| By KO | aspartate | Green | Green | Green | Green | Green | Green |
| By KO | glutamate | Green | Green | Green | Red | Green | Green |
| M00020 | serine | Red | Green | Green | Red | Green | Green |
| M00018 | threonine | Green | Green | Green | Red | Red | Green |
| By KO | asparagine | Green | Green | Green | Red | Green | Green |
| By KO | glutamine | Green | Green | Green | Red | Green | Green |
| M00021/M00338 | cysteine | Green | Yellow | Green | Red | Green | Green |
| By KO | glycine | Green | Green | Green | Green | Green | Green |
| M00015 | proline | Red | Red | Red | Red | Red | Green |
| By KO | alanine | Green | Green | Green | Red | Green | Green |
| M00019 | valine | Green | Yellow | Green | Red | Red | Green |
| M00019/M00570 | isoleucine | Green | Yellow | Green | Red | Red | Green |
| M00432+ KO | leucine | Green | Green | Green | Red | Red | Green |
| M00017 | methionine | Red | Yellow | Red | Red | Red | Green |
| M00024 | phenylalanine | Red | Red | Red | Red | Red | Yellow |
| M00025 | tyrosine | Red | Red | Red | Red | Red | Red |
| M00023 | tryptophan | Green | Green | Green | Red | Red | Green |

* 39% of MOα can synthesis serine and proline
** 30% of MOε can synthesis tyrosine
While MOα have few a.a. auxotrophies they have multiple a.a. transporters (as well as peptides). They also have an array of proteases. This indicates that they might first use external a.a. sources before spending energy on synthesizing their own.

B) Synthesis of vitamins/cofactors

| Module | Vitamin/Cofactor | MOα | MOγ | Brocadia | MOβ | MOδ | MOε |
|---|---|---|---|---|---|---|---|
| M00115 | NAD | Red | Green | Green | Red | Green | Green |
| M00116 | Menaquinone | Red | Red | Red | Red | Green | Red |
| M00117 | Ubiquinone | Red | Red | Red | Red | Red | Green |
| M00118 | Glutathione | Red | Red | Red | Red | Red | Green |
| M00120 | Coenzyme A | Green | Green | Green | Red | Green | Green |
| M00123 | Biotin (from pimeloyl) | Red | Red | Red | Red | Red | Green |
| M00124 | Pyridoxal | Red | Yellow | Green | Red | Green | Green |
| M00126 | Tetrahydrofolate | Red | Yellow | Red | Red | Green | Green |
| M00140 | C1-unit interconversion (bugs) | Green | Green | Green | Green | Green | Green |
| M00141 | C1-unit interconversion, (euks) | Green | Red | Red | Red | Red | Red |
| M00572 | Pimeloyl-ACP | Yellow | Green | Green | Red | Red | Green |
| M00573 | Biotin (from long chain acyl-ACP) | Red | Red | Red | Red | Red | Green |
| M00577 | Biotin (form pimelate) | Red | Red | Red | Red | Red | Green |
| By KO | Vitamin B12 | Red | Red | Green | Red | Red | Red |

* 39% of MOα can synthesize menaquinone
** 33% of MOγ can synthesize Biotin (per module) and menaquinone

B) Synthesis of lipids/fatty acids

| module | metabolite/process | MOα | MOγ | Brocadia | MOβ | MOδ | MOε |
|---|---|---|---|---|---|---|---|
| M00082 | F.a. initiation | Red | Red | Green | Red | Green | Green |
| M00083 | F.a. elongation | Red | Green | Red | Red | Green | Green |
| M00086 | acyl-CoA | Green | Green | Red | Red | Green | Green |
| M00087 | Beta oxidation | Green | Red | Red | Red | Red | Green |
| M00089 | Triacylglycerol | Red | Red | Red | Red | Red | Yellow |
| M00091 | Phosphatidylcholine | Red | Red | Green | Red | Red | Yellow |
| M00092 | Phosphatidylethanolamine (from PA) | Red | Red | Red | Red | Red | Red |
| M00098 | Acylglycerol | Red | Red | Red | Red | Red | Red |
| M00060 | KDO2-lipid A | Red | Green | Green | Red | Green | Green |
| M00063 | CMP-KDO | Red | Green | Green | Red | Green | Green |
| M00064 | ADP-L-glycero-D-manno-heptose | Red | Red | Green | Red | Green | Yellow |
| M00080 | O-antigen | Red | Red | Red | Red | Red | Red |

* nine MOα and six MOγ (all that are not complete) have 50% completeness for f.a initiation
** 33% of MOγ can initiate f.a. synthesis, synthesize phosphatidylcholine and ADP-L-glycero-D-manno-heptose, and have complete beta-oxidation
*** 30% of MOε can synthesize acylglycerol

#### C) Transport systems

| | Associated bacteria | Sludge associated bacteria | Anammox bacterium | CPR Bacteria | Ignavibacteriae/Bacteroides | Proteobacteria |
|---|---|---|---|---|---|---|
| metabolite | MOα | MOɣ | Brocadia | MOβ | MOδ | MOε |
| Capsular polysaccharide | 🔴 | 🔴 | 🔴 | 🔴 | 🔴 | 🔴 |
| Lipopolysaccharide | 🟢 | 🟢 | 🟢 | 🟢 | 🔴 | 🔴 |
| Lipooligosaccharide | 🔴 | 🔴 | 🔴 | 🔴 | 🔴 | 🔴 |
| Sodium | 🔴 | 🔴 | 🔴 | 🔴 | 🔴 | 🔴 |
| ABC-2 type | 🟢 | 🟢 | 🟢 | 🟢 | 🟢 | 🟢 |
| Lipoprotein-releasing system | 🔴 | 🔴 | 🔴 | 🔴 | 🟢 | 🟢 |
| Cell division | 🔴 | 🟡 | 🔴 | 🟢 | 🟢 | 🟢 |
| Putative ABC | 🟢 | 🟢 | 🟢 | 🔴 | 🟢 | 🟢 |
| Heme | 🟢 | 🟢 | 🔴 | 🔴 | 🟢 | 🟢 |
| Lipopolysaccharide export system | 🔴 | 🟢 | 🟢 | 🔴 | 🟢 | 🟢 |
| Iron complex | 🟢 | 🟢 | 🟢 | 🔴 | 🔴 | 🟢 |
| Vitamin B12 | 🔴 | 🔴 | 🔴 | 🔴 | 🔴 | 🔴 |
| Zinc | 🔴 | 🔴 | 🔴 | 🔴 | 🟢 | 🔴 |
| Manganese/iron | 🟢 | 🔴 | 🔴 | 🔴 | 🔴 | 🔴 |
| Putative zinc/manganese | 🔴 | 🔴 | 🟢 | 🔴 | 🔴 | 🔴 |
| Cobalt/nickel | 🔴 | 🔴 | 🟢 | 🔴 | 🔴 | 🔴 |
| Nickel | 🔴 | 🔴 | 🟢 | 🔴 | 🔴 | 🔴 |
| Manganese/zinc/iron | 🔴 | 🔴 | 🔴 | 🔴 | 🔴 | 🔴 |
| Biotin | 🔴 | 🔴 | 🔴 | 🔴 | 🔴 | 🔴 |
| Energy-coupling factor | 🟢 | 🔴 | 🔴 | 🔴 | 🔴 | 🔴 |
| Sulfate | 🔴 | 🔴 | 🔴 | 🔴 | 🔴 | 🔴 |
| Tungstate | 🟢 | 🔴 | 🔴 | 🔴 | 🔴 | 🟡 |
| NitT/TauT family | 🟢 | 🟢 | 🟢 | 🔴 | 🔴 | 🟢 |
| Molybdate | 🟢 | 🔴 | 🟢 | 🔴 | 🔴 | 🟢 |
| Iron(III) | 🟢 | 🔴 | 🔴 | 🔴 | 🔴 | 🟢 |
| Thiamine | 🟢 | 🔴 | 🔴 | 🔴 | 🔴 | 🔴 |
| Putative spermidine/putrescine | 🟢 | 🔴 | 🔴 | 🔴 | 🔴 | 🔴 |
| Glycine betaine/proline | 🔴 | 🔴 | 🔴 | 🔴 | 🔴 | 🔴 |
| Osmoprotectant | 🔴 | 🔴 | 🔴 | 🔴 | 🔴 | 🔴 |
| Spermidine/putrescine | 🟢 | 🔴 | 🔴 | 🔴 | 🔴 | 🔴 |
| Putrescine | 🔴 | 🔴 | 🔴 | 🔴 | 🔴 | 🟡 |
| Peptides/nickel | 🟢 | 🟢 | 🔴 | 🔴 | 🟢 | 🟢 |
| Microcin C | 🔴 | 🔴 | 🔴 | 🔴 | 🔴 | 🔴 |
| Oligopeptide | 🔴 | 🔴 | 🔴 | 🔴 | 🔴 | 🔴 |
| Phosphate | 🟢 | 🔴 | 🟢 | 🔴 | 🔴 | 🟢 |
| Phosphonate | 🟢 | 🔴 | 🔴 | 🔴 | 🔴 | 🔴 |
| Glutamate/aspartate | 🔴 | 🔴 | 🔴 | 🔴 | 🔴 | 🔴 |
| General L-amino acid | 🔴 | 🔴 | 🔴 | 🔴 | 🔴 | 🔴 |
| Putative polar amino acid | 🟢 | 🔴 | 🔴 | 🔴 | 🔴 | 🔴 |
| Branched-chain amino acid | 🟢 | 🔴 | 🔴 | 🔴 | 🔴 | 🟢 |
| Raffinose/stachyose/melibiose | 🟢 | 🔴 | 🔴 | 🔴 | 🔴 | 🔴 |
| Putative sn-glycerol-phosphate | 🔴 | 🔴 | 🔴 | 🔴 | 🔴 | 🔴 |
| alpha-Glucoside | 🟢 | 🔴 | 🔴 | 🔴 | 🔴 | 🔴 |
| Trehalose/maltose | 🔴 | 🔴 | 🔴 | 🔴 | 🔴 | 🔴 |
| Putative multiple sugar | 🟢 | 🟢 | 🔴 | 🔴 | 🔴 | 🔴 |
| Phospholipid | 🔴 | 🔴 | 🟢 | 🔴 | 🔴 | 🟢 |
| Ribose | 🟢 | 🔴 | 🔴 | 🔴 | 🔴 | 🔴 |
| D-Xylose | 🔴 | 🔴 | 🔴 | 🔴 | 🔴 | 🔴 |
| Multiple sugar | 🔴 | 🔴 | 🔴 | 🔴 | 🔴 | 🔴 |
| Putative simple sugar | 🟢 | 🔴 | 🔴 | 🔴 | 🔴 | 🔴 |
| arabinogalactan oligomer/maltooligosaccharide | 🟢 | 🔴 | 🔴 | 🔴 | 🔴 | 🔴 |
| Glucose/mannose | 🔴 | 🔴 | 🔴 | 🔴 | 🔴 | 🔴 |
| gamma-Hexachlorocyclohexane | 🔴 | 🟢 | 🟢 | 🔴 | 🟢 | 🟢 |
| Mce | 🔴 | 🟢 | 🟢 | 🔴 | 🟢 | 🟢 |

**Table S6** Confidence intervals of the Log-Ratio changes for each reference frame genome (RFg)

| | Upper Limit | Lower Limit |
|---|---|---|
| RF1 (LAC_NA06_Anaerolineales_42_27) | -0.07005769 | -0.7292107 |
| RF2 (anamox4_Bacteria_63_7_curated) | -0.07005769 | -0.7292107 |
| RF3 (anamox4_Gammaproteobacteria_67_14_curated) | 0.3940631 | -0.390227 |
| RF4 (LAC_NA07_Proteobacteria_68_32) | -0.2433609 | -0.8117102 |

**Table S7** Closest relatives based on distance in ML tree, for the anammox bacterial community

| **Genome** | **Closest relative** | **Distance** | **Similarity** |
|---|---|---|---|
| **LAC_ACD01** | **Actinobacteria_bacterium_RBG_16_70_17** | **0.58** | **88.56** |
| **LAC_ACD02** | **LAC_ACD07** | **0.42** | **91.74** |
| **LAC_BAC01** | **OLB21** | **0.78** | **84.69** |
| **LAC_BAC02** | **LAC_BAC15** | **0.59** | **88.39** |
| **LAC_BAC03** | **CG10_big_fil_rev_8_21_14_0_10_Saccharibacteria_47_8** | **1.17** | **76.89** |
| **LAC_BAC04** | **Deltaproteobacteria_bacterium_GWA2_45_12** | **0.88** | **82.55** |
| **LAC_BAC05** | **OLB16** | **0.00** | **99.92** |
| **LAC_CHLX01** | **UTCFX4** | **0.12** | **97.66** |
| **LAC_BAC06** | **Bacteria_Lentisphaerae_GWF2_Lentisphaerae_57_35** | **0.66** | **87.00** |
| **LAC_BAC07** | **LAC_CHLX01** | **0.25** | **95.01** |
| **LAC_BAC08** | **LAC_BAC09** | **0.31** | **93.82** |
| **LAC_BAC09** | **LAC_BAC08** | **0.31** | **93.82** |
| **LAC_BAC10** | **Bacteria_Plantomycetes_uncultured_DG_23** | **1.36** | **73.17** |
| **LAC_BAC11** | **Bacteria_Acidobacteria_RBG_16_Acidobacteria_64_8** | **0.34** | **93.22** |
| **LAC_BACT01** | **Bacteria_BacteroidetesChlorobi_group_Bacteroidetes_Bacteroidetes_Order_III__Incertae_sedis_Thermonema_rossianum_DSM_10300** | **0.75** | **85.21** |
| **LAC_BACT02** | **OLB10** | **0.01** | **99.75** |
| **LAC_BACT03** | **Bacteria_BacteroidetesChlorobi_group_Bacteroidetes_Flavobacteriia_Flavobacteriales_Cryomorphaceae_Crocinitomix_catalasitica_ATCC_23190** | **0.68** | **86.51** |
| **LAC_PROT01** | **CG18_big_fil_WC_8_21_14_2_50_Hydrogenophilales_58_12** | **0.54** | **89.32** |
| **LAC_PROT02** | **LAC_PROT03** | **0.08** | **98.33** |
| **LAC_PROT03** | **LAC_PROT02** | **0.08** | **98.33** |
| **LAC_CLO01** | **Bacteria_CP_WWE1_candidate_division_WWE1_bacterium_JGI_0000039_M09_TAsludge_001_159** | **0.01** | **99.86** |

| | | | |
|---|---|---|---|
| **LAC_CHLX02** | **UTCFX1** | **0.00** | **100.00** |
| **LAC_PROT04** | **Bacteria_Proteobacteria_Gammaproteobacteria_Alteromonadales_Alteromonadaceae_Melitea_salexigens_DSM_19753** | **0.43** | **91.44** |
| **LAC_GMT01** | **Bacteria_Gemmatimonadetes_Gemmatimonadetes_Gemmatimonadales_Gemmatimonadaceae_Gemmatimonas_aurantiaca_T_27** | **0.12** | **97.58** |
| **LAC_PROT05** | **LAC_PROT17** | **0.03** | **99.36** |
| **LAC_PROT06** | **LAC_PROT19** | **0.22** | **95.59** |
| **LAC_PLT01** | **Bacteria_Planctomycetes_Planctomycetia_Planctomycetales_Planctomycetaceae_Pirellula_staleyi_DSM_6068** | **0.61** | **88.05** |
| **LAC_PROT07** | **LAC_PROT20** | **0.08** | **98.41** |
| **LAC_PROT08** | **LAC_PROT20** | **0.27** | **94.68** |
| **LAC_ACD03** | **OLB17** | **0.18** | **96.39** |
| **LAC_IGN01** | **Bacteria_Ignavibacteria_RBG_16_Ignavibacteria_36_9** | **0.10** | **97.97** |
| **LAC_PROT09** | **UTPRO2** | **0.13** | **97.49** |
| **LAC_BACT04** | **UTBCD1** | **0.05** | **99.03** |
| **LAC_BACT05** | **Sphingobacteriia_bacterium_RIFOXYD2_FULL_35_12** | **0.35** | **93.06** |
| **LAC_D-T01** | **LAC_D-T02** | **0.01** | **99.87** |
| **LAC_SPR01** | **Bacteria_Spirochaetes_Spirochaetia_Spirochaetales_Leptospiraceae_Turneriella_parva_H_DSM_21527** | **0.25** | **95.08** |
| **LAC_ACT01** | **Bacteria_Actinobacteria_Actinobacteria_unclassified_Actinobacteria_Candidatus_Microthrix_parvicella_Bio17_1** | **0.41** | **91.84** |
| **LAC_BAC12** | **OLB21** | **1.22** | **75.94** |
| **LAC_BAC13** | **Bacteria_Armatimonadetes_Fimbriimonas_ginsengisoli_Gsoil_348** | **0.37** | **92.70** |
| **LAC_BAC14** | **Bacteria_Cyanobacteria_Oscillatoriales_Microcoleus_sp__PCC_7113** | **1.00** | **80.31** |
| **LAC_BACT06** | **CG18_big_fil_WC_8_21_14_2_50_Bacteroidetes_41_14** | **0.40** | **92.02** |
| **LAC_PROT10** | **LAC_PROT27** | **0.16** | **96.82** |
| **LAC_PROT11** | **Burkholderiales_bacterium_RIFCSPLOWO2_12_FULL_65_40** | **0.08** | **98.46** |
| **LAC_PROT12** | **LAC_PROT27** | **0.22** | **95.59** |
| **LAC_CHLX03** | **LAC_CHLX09** | **0.13** | **97.47** |
| **LAC_PROT13** | **Bacteria_Proteobacteria_Gammaproteobacteria_Thiothrichales_uncultured_SG8_50** | **0.28** | **94.54** |
| **LAC_PROT14** | **Bacteria_Proteobacteria_Betaproteobacteria_Hydrogenophilales_GWE1_Thiobacillus_62_9** | **0.10** | **98.12** |
| **LAC_IGN02** | **LAC_IGN05** | **0.03** | **99.39** |
| **LAC_IGN03** | **UTCHB2** | **0.16** | **96.84** |
| **LAC_IGN04** | **Bacteria_Ignavibacteria_GWA2_Ignavibacteriae_55_25** | **0.52** | **89.66** |
| **LAC_ARCH01** | **Archaea_Euryarchaeota_Methanomicrobia_Methanosarcinales_Methanosarcinaceae_Methanosarcina_barkeri_Fusaro_DSM_804** | **0.10** | **97.99** |
| **LAC_MIC01** | **Candidatus_Collierbacteria_bacterium_RIFOXYB1_FULL_49_13** | **0.75** | **85.11** |

| | | | |
|---|---|---|---|
| **LAC_MIC02** | **LAC_MIC04** | **0.59** | **88.44** |
| **LAC_PROT15** | **Bacteria_Proteobacteria_deltaepsilon_subdivisions_Deltaproteobacteria_Myxococcales_Sorangiineae_Polyangiaceae_Chondromyces_apiculatus_DSM_436** | **0.35** | **93.10** |
| **LAC_PROT16** | **Bacteria_Proteobacteria_Alphaproteobacteria_Rhizobiales_Phyllobacteriaceae_Aquamicrobium_defluvii_W13Z1** | **0.08** | **98.32** |
| **LAC_MIC03** | **UTCPR1** | **0.00** | **100.00** |
| **LAC_BACT07** | **UTBCD1** | **0.05** | **98.99** |
| **LAC_IGN05** | **LAC_IGN02** | **0.03** | **99.39** |
| **LAC_VER01** | **LAC_VER02** | **0.09** | **98.18** |
| **LAC_VER02** | **LAC_VER01** | **0.09** | **98.18** |
| **LAC_PROT17** | **LAC_PROT05** | **0.03** | **99.36** |
| **LAC_PROT18** | **Bacteria_Proteobacteria_Gammaproteobacteria_Xanthomonadales_Xanthomonadaceae_Rudaea_cellulosilytica_DSM_22992** | **0.28** | **94.43** |
| **LAC_ACD04** | **Bacteria_Acidobacteria_RIFCSPLOWO2_02_FULL_Acidobacteria_68_18** | **0.40** | **92.19** |
| **LAC_BAC15** | **OLB6** | **0.00** | **99.96** |
| **LAC_BAC16** | **Bacteria_ChlamydiaeVerrucomicrobia_group_Verrucomicrobia_Spartobacteria_Chthoniobacter_flavus_Ellin428_unfinished_sequence** | **0.95** | **81.20** |
| **LAC_BAC18** | **Bacteria_Chloroflexi_RBG_16_Chloroflexi_68_14** | **0.74** | **85.42** |
| **LAC_BACT08** | **LAC_BACT13** | **0.66** | **87.00** |
| **LAC_IGN06** | **Bacteria_RIF-IGX_RIFOXYC2_FULL_RIF_IGX_35_21** | **0.59** | **88.31** |
| **LAC_BACT09** | **LAC_PROT25** | **0.00** | **100.00** |
| **LAC_CHLX04** | **LAC_CHLX08** | **0.11** | **97.80** |
| **LAC_CHLX05** | **CG2_30_FULL_Chloroflexi_64_16** | **0.32** | **93.76** |
| **LAC_PROT19** | **OLB2** | **0.00** | **99.99** |
| **LAC_OMN01** | **Bacteria_OP3X_RIFCSPHIGHO2_02_FULL_OP3X_63_14** | **0.49** | **90.34** |
| **LAC_VER03** | **Bacteria_ChlamydiaeVerrucomicrobia_group_Verrucomicrobia_Opitutae_Opitutales_Opitutaceae_Opitutus_terrae_PB90_1** | **0.18** | **96.48** |
| **LAC_VER04** | **Bacteria_ChlamydiaeVerrucomicrobia_group_Verrucomicrobia_Verrucomicrobiae_Verrucomicrobiales_Verrucomicrobia_subdivision_3_bacterium_Ellin514** | **0.46** | **90.90** |
| **LAC_PROT20** | **LAC_PROT07** | **0.08** | **98.41** |
| **LAC_BACT10** | **Bacteria_BacteroidetesChlorobi_group_Bacteroidetes_Sphingobacteriia_Sphingobacteriales_Chitinophagaceae_Niastella_koreensis_GR20_10_DSM_17620** | **0.31** | **93.92** |
| **LAC_BACT11** | **Bacteria_BacteroidetesChlorobi_group_Bacteroidetes_Sphingobacteriia_Sphingobacteriales_Saprospiraceae_Haliscomenobacter_hydrossis_O_DSM_1100** | **0.48** | **90.51** |
| **LAC_IGN07** | **UTCHB3** | **0.00** | **100.00** |
| **LAC_VER05** | **Bacteria_ChlamydiaeVerrucomicrobia_group_Verrucomicrobia_Spartobacteria_Chthoniobacter_flavus_Ellin428_unfinished_sequence** | **0.53** | **89.54** |

| | | | |
|---|---|---|---|
| LAC_PROT21 | Bacteria_Proteobacteria_Alphaproteobacteria_unclassified_Alphaproteobacteria_Micavibrio_aeruginosavorus_ARL_13 | 0.65 | 87.09 |
| LAC_PROT22 | OLB13 | 0.00 | 99.99 |
| LAC_BAC20 | UTPRO1 | 0.00 | 100.00 |
| LAC_BACT12 | LAC_BACT02 | 0.40 | 92.12 |
| LAC_NIT01 | OLB3 | 0.00 | 99.94 |
| LAC_CHLX06 | CG2_30_FULL_Chloroflexi_64_16 | 0.63 | 87.65 |
| LAC_PROT22 | Gammaproteobacteria_bacterium_RIFCSPLOWO2_12_FULL_52_10 | 0.64 | 87.35 |
| LAC_MIC04 | LAC_MIC02 | 0.59 | 88.44 |
| LAC_PROT23 | LAC_PROT19 | 0.24 | 95.17 |
| LAC_PLT02 | UTAMX1 | 0.00 | 100.00 |
| LAC_CHLX07 | Bacteria_Chloroflexi_GWB2_Chloroflexi_54_36 | 0.40 | 92.09 |
| LAC_BAC21 | LAC_BAC20 | 0.61 | 88.00 |
| LAC_BACT13 | LAC_BACT08 | 0.66 | 87.00 |
| LAC_PROT24 | Betaproteobacteria_bacterium_RIFCSPLOWO2_12_FULL_62_13b | 0.52 | 89.79 |
| LAC_PROT25 | LAC_BACT09 | 0.00 | 100.00 |
| LAC_SCH01 | CG_4_10_14_0_2_um_filter_Saccharibacteria_52_9 | 0.69 | 86.30 |
| LAC_ACD05 | LAC_ACD07 | 0.38 | 92.40 |
| LAC_CHLX08 | UTCFX2 | 0.00 | 99.99 |
| LAC_ARM01 | OLB18 | 0.00 | 99.95 |
| LAC_MIC05 | LAC_MIC04 | 0.81 | 83.95 |
| LAC_PROT26 | Bacteria_Proteobacteria_Alphaproteobacteria_Rhodobacterales_Rhodobacteraceae_Rhodovulum_sp__PH10 | 0.28 | 94.38 |
| LAC_DADA01 | Candidatus_Dadabacteria_bacterium_RIFCSPHIGHO2_12_FULL_53_21 | 0.25 | 95.11 |
| LAC_CHLX09 | LAC_CHLX03 | 0.13 | 97.47 |
| LAC_CHLX10 | UTCFX3 | 0.00 | 99.93 |
| LAC_ACD06 | Bacteria_Acidobacteria_RBG_13_Acidobacteria_68_16 | 0.33 | 93.48 |
| LAC_ACT02 | Actinobacteria_bacterium_RBG_16_68_12 | 0.31 | 93.96 |
| LAC_ACT03 | LAC_ACT01 | 0.58 | 88.52 |
| LAC_ACT04 | Bacteria_Actinobacteria_Actinobacteria_Actinobacteridae_Actinomycetales_Micrococcineae_Cellulomonadaceae_Actinotalea_ferrariae_CF5_4 | 0.17 | 96.58 |
| LAC_BAC22 | UTCHB1 | 0.00 | 100.00 |
| LAC_BAC23 | LAC_BAC08 | 0.84 | 83.47 |
| LAC_BAC24 | Bacteria_RIF-CHLX_GWC2_RIF_CHLX_73_18 | 0.40 | 92.09 |
| LAC_PROT27 | LAC_PROT10 | 0.16 | 96.82 |
| LAC_CHLX11 | Bacteria_Chloroflexi_Caldilineae_Caldilineales_Caldilineaceae_Caldilinea_a | 0.22 | 95.58 |

| | | | |
|---|---|---|---|
| | erophila_STL_6_O1_DSM_14535 | | |
| LAC_ACD07 | LAC_ACD05 | 0.38 | 92.40 |
| LAC_CHLX12 | LAC_CHLX06 | 0.82 | 83.78 |
| LAC_CHLX13 | Bacteria_Chloroflexi_Anaerolineae_Anaerolineales_Anaerolineaceae_Anaerolinea_thermophila_UNI_1 | 0.28 | 94.43 |
| LAC_CHLX14 | OLB15 | 0.37 | 92.70 |
| LAC_CHLX15 | LAC_CHLX17 | 0.64 | 87.28 |
| LAC_CHLX16 | LAC_CHLX17 | 0.25 | 94.97 |
| LAC_CHLX17 | LAC_CHLX16 | 0.25 | 94.97 |
| LAC_PROT28 | LAC_PROT08 | 0.59 | 88.31 |
| LAC_CHLX18 | Bacteria_RIF-CHLX_RBG_16_RIF_CHLX_72_14 | 0.14 | 97.17 |
| LAC_PROT29 | Rhodobacteraceae_bacterium_GWF1_65_7 | 0.28 | 94.38 |
| LAC_PROT30 | UTPRO2 | 0.00 | 99.97 |
| LAC_MIC06 | Candidatus_Roizmanbacteria_bacterium_RIFOXYA1_FULL_41_12 | 0.70 | 86.25 |
| LAC_D-T02 | LAC_D-T01 | 0.01 | 99.87 |
| OLB1 | Candidatus_Brocadia_sinica_JPN1 | 0.00 | 99.90 |
| OLB10 | LAC_BACT02 | 0.01 | 99.75 |
| OLB11 | Bacteria_BacteroidetesChlorobi_group_Bacteroidetes_Sphingobacteriia_Sphingobacteriales_Chitinophagaceae_Chitinophaga_pinensis_DSM_2588 | 0.41 | 91.93 |
| OLB12 | Bacteria_BacteroidetesChlorobi_group_Bacteroidetes_Cytophagia_Cytophagales_Flammeovirgaceae_Fulvivirga_imtechensis_AK7 | 0.39 | 92.33 |
| OLB13 | LAC_PROT22 | 0.00 | 99.99 |
| OLB14 | Bacteria_Chloroflexi_RBG_16_Chloroflexi_51_16 | 0.32 | 93.67 |
| OLB15 | LAC_CHLX14 | 0.37 | 92.70 |
| OLB16 | LAC_BAC05 | 0.00 | 99.92 |
| OLB17 | LAC_ACD03 | 0.18 | 96.39 |
| OLB18 | LAC_ARM01 | 0.00 | 99.95 |
| OLB2 | LAC_PROT19 | 0.00 | 99.99 |
| OLB20 | OLB21 | 0.41 | 91.82 |
| OLB21 | OLB20 | 0.41 | 91.82 |
| OLB22 | Candidatus_Roizmanbacteria_bacterium_RIFCSPHIGHO2_01_FULL_39_12b | 1.04 | 79.49 |
| OLB23 | Candidatus_Roizmanbacteria_bacterium_RIFCSPLOWO2_01_FULL_38_11 | 0.62 | 87.77 |
| OLB3 | LAC_NIT01 | 0.00 | 99.94 |
| OLB4 | LAC_BAC22 | 0.00 | 99.99 |
| OLB5 | UTCHB1 | 0.62 | 87.78 |

| | | | |
|---|---|---|---|
| **OLB6** | **LAC_BAC15** | **0.00** | **99.96** |
| **OLB7** | **Bacteria_RIF-IGX_RIFOXYC2_FULL_RIF_IGX_35_21** | **0.87** | **82.83** |
| **OLB8** | **OLB9** | **0.58** | **88.57** |
| **OLB9** | **OLB8** | **0.58** | **88.57** |
| **UTAMX1** | **LAC_PLT02** | **0.00** | **100.00** |
| **UTAMX2** | **UTAMX1** | **0.16** | **96.76** |
| **UTBCD1** | **LAC_BACT04** | **0.05** | **99.03** |
| **UTCFX1** | **LAC_CHLX02** | **0.00** | **100.00** |
| **UTCFX2** | **LAC_CHLX08** | **0.00** | **99.99** |
| **UTCFX3** | **LAC_CHLX10** | **0.00** | **99.93** |
| **UTCFX4** | **LAC_CHLX01** | **0.12** | **97.66** |
| **UTCFX5** | **LAC_PROT22** | **0.00** | **99.99** |
| **UTCHB1** | **LAC_BAC22** | **0.00** | **100.00** |
| **UTCHB2** | **LAC_IGN05** | **0.05** | **99.10** |
| **UTCHB3** | **LAC_IGN07** | **0.00** | **100.00** |
| **UTCPR1** | **LAC_MIC03** | **0.00** | **100.00** |
| **UTPLA1** | **Bacteria_Plantomycetes_Phycisphaerae_uncultured_SMTZ_30** | **0.95** | **81.25** |
| **UTPRO1** | **LAC_BAC20** | **0.00** | **100.00** |
| **UTPRO2** | **LAC_PROT30** | **0.00** | **99.97** |

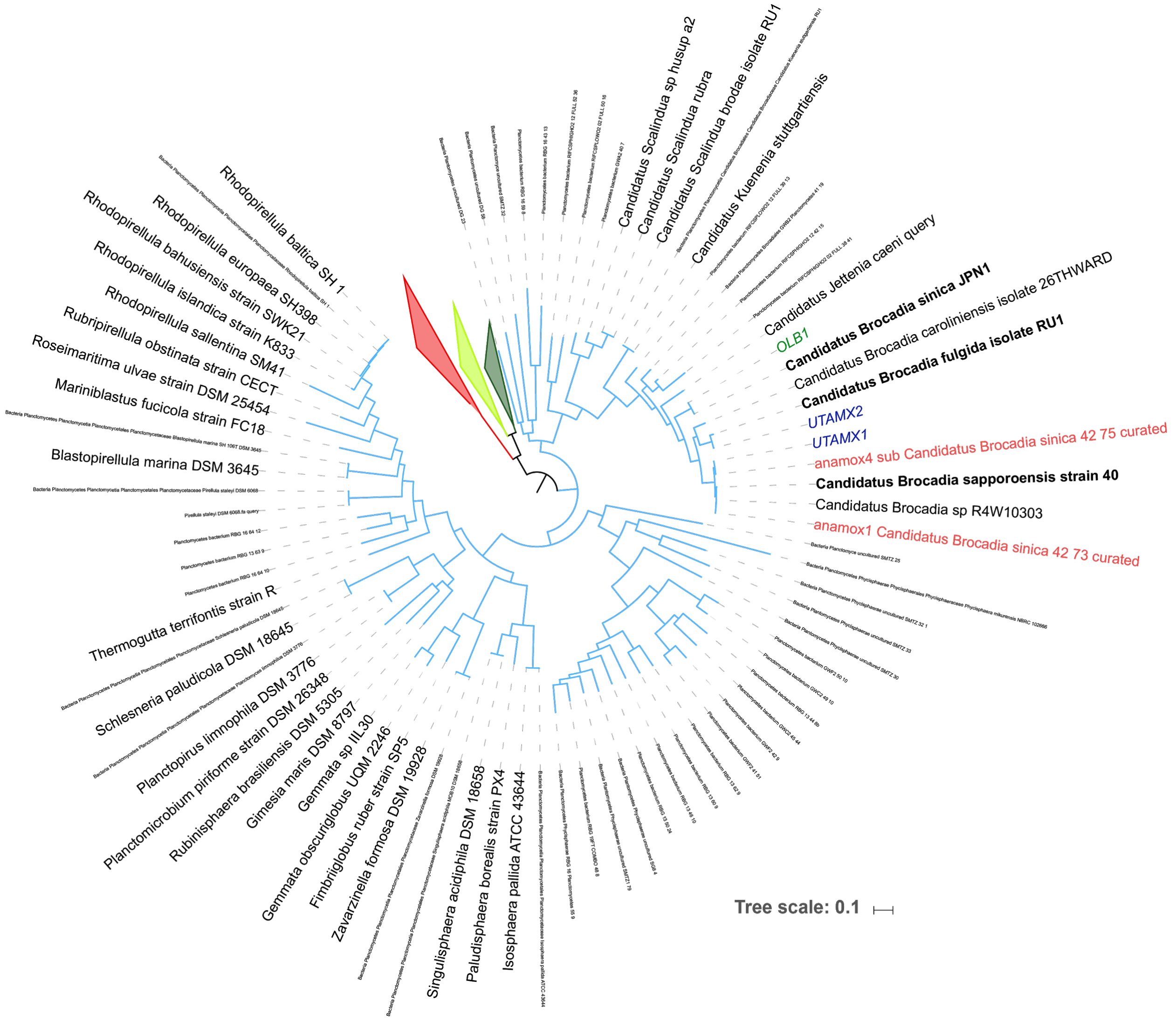

**Figure S3 Maximum likelihood tree of the PVC clade (*Planctomycetes*, *Verrucomicrobia*, *Chlamydiae*, and *Lentisphaerae*).** This tree is based on amino acid sequence alignment of 15 concatenated Ribosomal Proteins. Genomes from the current study, genomes from previous studies, and all reference sequences were used for the construction of the tree. To these sequences, 35 *Planctomycetes* were added from NCBI, bringing the total number of sequences to 123. Sequences were aligned with MAFFT (default parameters) and a RaxML tree was constructed in The CIPRES Science Gateway V. 3.3. Branch colors follow the ggkbase phyla color palette. The *Chlamydiae*, *Verrucomicrobia*, and *Lentisphaerae* phyla are collapsed to allow easier viewing of the *Planctomycetes*. Genomes of anammox bacteria from the current paper and previous papers are colored according to marking in Figure 2A. *Planctomycetes* genomes from NCBI are in larger font, and closest NCBI references to the anammox bioreactor's genomes are in bold. All anammox bacteria are of the genus *Brocadia*. The anammox bacterium from the current study (both strains are presented) and the dominant anammox bacterium from Lawson et al. are both closely related to *Brocadia sapporoensis*. The second anammox bacterium from the Lawson et al. study is closely related to *Brocadia fulgida*. The anammox bacterium from the Speth et al. study is closely related to *Brocadia sinica*
.

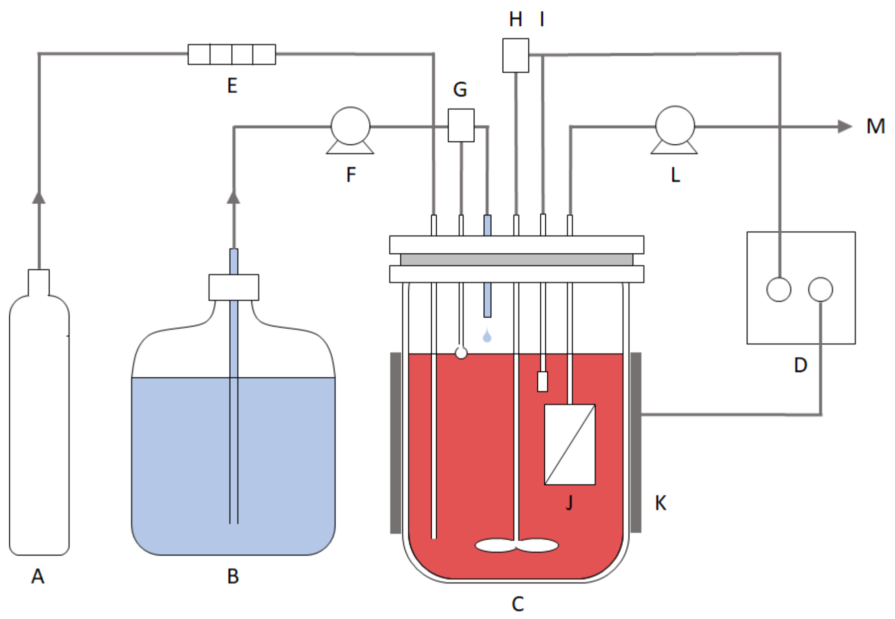

**Figure S4 Schematic of the anaerobic membrane bioreactor.** The components of the anaerobic membrane bioreactor include: A, influent gas tank; B, influent media tank; C, 1L reactor vessel; D, power supply device; E, flowmeter; F, influent peristaltic pump; G, level controller; H, impeller; I, temperature probe; J, membrane module; K, heating jacket; L, effluent peristaltic pump; and M, effluent line.

**Table S8 Composition of the synthetic wastewater feed.** This recipe, based on van de Graaf et al. 1995, was modified to better mimic the composition of the anaerobic digester sidestream at East Bay Municipal Utility District (Oakland, CA). Concentrations of nitrogen species varied with time, (Figure 3). Additionally, the concentrations of the four constituents asterisked below--iron, zinc, molybdenum, and copper--were increased from their lower to higher values on Day 353.

| Constituent | Concentration | Unit |
|---|---|---|
| $(NH_4)_2SO_4$ | 40-500 | mg-N/L |
| $NaNO_2$ | 5-660 | mg-N/L |
| NaCl | 1000 | mg/L |
| $MgCl_2 \cdot 6H_2O$ | 500 | mg/L |
| $KH_2PO_4$ | 200 | mg/L |
| KCl | 300 | mg/L |
| $CaCl_2 \cdot 2H_2O$ | 180 | mg/L |
| $KHCO_3$ | 420 | mg/L |
| *$FeCl_2 \cdot 4H_2O$ | 7.7; 17.9 | mg/L |
| $CoCl_2 \cdot 6H_2O$ | 0.24 | mg/L |
| $MnCl_2 \cdot 4H_2O$ | 0.99 | mg/L |
| *$ZnCl_2$ | 0.07; 0.20 | mg/L |
| $H_3BO_3$ | 0.014 | mg/L |
| *$Na_2MoO_4 \cdot 2H_2O$ | 0.10; 0.22 | mg/L |
| $NiCl_2 \cdot 6H_2O$ | 0.19 | mg/L |
| *$CuCl_2 \cdot 2H_2O$ | 0.002; 0.17 | mg/L |
| $Na_2SeO_3 \cdot 5H_2O$ | 0.16 | mg/L |
| pH | 6.8-7.0 | - |

### Supplemental methods

#### Refinement of the *Candidatus Brocadia* sp. genome.

After running dRep but before running ra2, we discovered that the hydrazine synthase (Hzs) gene, fundamental to the anammox process, was not binned with the *Brocadia* sp. genome. The Hzs gene was skewed in both GC and CV, but the taxonomic identification of the scaffold linked it to the *Brocadia* sp. genome. No other Planctomycetes genomes were found in the GC and CV ranges that covered the Hzs-harboring scaffold.

To test if the Hzs gene belonged to the *Brocadia* sp. genome, we clustered scaffolds according to their tetramer frequency based on Emergent Self-Organizing Maps (ESOMs) with Databionic ESOM Analyzer [59]. In summary, we downloaded the *Brocadia* sp. genome, a temporary bin with the Hzs gene, and a second temporary bin with all other *Planctomycetes* scaffolds within the GC and CV ranges of both prior bins. To each of these bins we added representative genomes with varying CV values, and also 12 *Brocadiales* genomes found in NCBI Genome database. The *Brocadia* sp. genome, along with the two temporary bins, clustered with *Brocadia sinica*. Following this analysis, both the Hzs bin and the tmp bin were added to the *Brocadia* sp. bin.

**Table S9** Replication rates of genomes for each time point, calculated using the iRep program.

| Genome | D0 | D82 | D166 | D284 | D328 | D437 |
|---|---|---|---|---|---|---|
| LAC_BAC01 | 1.46 | NA | 1.44 | NA | NA | NA |
| LAC_BAC02 | 1.40 | NA | NA | NA | NA | NA |
| LAC_BAC03 | 1.30 | NA | 1.42 | NA | NA | NA |
| LAC_BAC04 | 1.21 | NA | 1.30 | NA | NA | NA |
| LAC_BAC05 | 1.34 | NA | 1.40 | NA | NA | NA |
| LAC_CHLX01 | 1.31 | 1.40 | 1.16 | 1.32 | 1.11 | 1.21 |
| LAC_BAC06 | 1.25 | NA | 1.28 | NA | NA | NA |
| LAC_BAC08 | 1.31 | NA | 1.38 | NA | NA | NA |
| LAC_BAC09 | 1.36 | NA | 1.40 | NA | NA | NA |
| LAC_BAC11 | 1.28 | NA | 1.30 | NA | NA | NA |
| LAC_BACT02 | 1.32 | 1.44 | 1.39 | NA | NA | NA |
| LAC_BACT03 | 1.30 | 1.21 | 1.33 | NA | 1.37 | NA |
| LAC_PROT02 | 1.15 | 1.18 | 1.17 | 1.30 | 1.30 | NA |
| LAC_PROT03 | 1.27 | 1.31 | 1.32 | 1.40 | 1.36 | NA |
| LAC_CHLX02 | 1.31 | 1.31 | 1.16 | 1.17 | 1.43 | NA |
| LAC_PROT04 | 1.42 | NA | NA | NA | NA | NA |
| LAC_GMT01 | 1.34 | NA | 1.39 | NA | NA | NA |
| LAC_PROT05 | 1.67 | NA | NA | NA | NA | NA |
| LAC_PROT06 | 1.24 | NA | 1.29 | NA | NA | NA |
| LAC_PROT07 | 1.55 | NA | 1.62 | NA | NA | NA |
| LAC_PROT08 | 1.45 | NA | 1.53 | NA | NA | NA |
| LAC_ACD03 | 1.36 | 1.13 | 1.19 | 1.35 | 1.29 | NA |
| LAC_PROT09 | 1.41 | NA | NA | NA | NA | NA |
| LAC_BACT04 | 1.31 | 1.32 | 1.26 | 1.26 | 1.18 | 1.28 |
| LAC_BACT05 | 1.41 | NA | NA | NA | NA | NA |
| LAC_SPR01 | 1.37 | NA | NA | NA | NA | NA |
| LAC_BAC13 | NA | 1.35 | NA | NA | NA | NA |
| LAC_BACT06 | NA | 1.37 | NA | NA | NA | NA |
| LAC_PROT11 | NA | 1.35 | NA | NA | NA | NA |
| LAC_PROT12 | NA | 1.29 | NA | NA | NA | NA |
| LAC_CHLX03 | NA | 1.36 | NA | NA | NA | NA |
| LAC_PROT13 | NA | 1.28 | NA | NA | NA | NA |
| LAC_PROT14 | NA | 1.32 | NA | NA | NA | NA |
| LAC_IGN02 | NA | 1.27 | 1.81 | NA | NA | NA |
| LAC_IGN03 | NA | 1.46 | NA | NA | NA | NA |
| LAC_IGN04 | NA | 1.29 | NA | NA | NA | NA |
| LAC_ARCH01 | NA | 1.31 | NA | NA | NA | NA |

| | | | | | | |
|---|---|---|---|---|---|---|
| LAC_MIC01 | NA | 1.47 | NA | NA | 1.42 | 1.36 |
| LAC_MIC02 | NA | 1.39 | NA | NA | 2.57 | 2.40 |
| LAC_PROT16 | NA | 1.17 | NA | NA | NA | NA |
| LAC_MIC03 | NA | 1.54 | 1.60 | NA | NA | NA |
| LAC_BACT07 | NA | 1.31 | NA | NA | NA | NA |
| LAC_IGN05 | 1.42 | 1.31 | 1.38 | 1.51 | 1.58 | 1.90 |
| LAC_VER01 | NA | 1.38 | NA | NA | NA | NA |
| LAC_VER02 | 1.34 | 1.37 | 1.38 | NA | 1.44 | NA |
| LAC_PROT17 | NA | 1.28 | NA | NA | NA | NA |
| LAC_PROT18 | NA | 1.51 | NA | NA | 1.56 | NA |
| LAC_BAC16 | NA | NA | 1.36 | NA | NA | NA |
| LAC_BAC17 | NA | NA | 1.25 | NA | NA | NA |
| LAC_BAC18 | 1.37 | NA | 1.34 | NA | NA | NA |
| LAC_BACT08 | 1.22 | NA | 1.30 | NA | NA | NA |
| LAC_IGN06 | NA | 1.41 | 1.33 | NA | 1.36 | NA |
| LAC_PROT19 | 1.17 | 1.38 | 1.22 | 1.32 | 1.38 | NA |
| LAC_OMN01 | NA | NA | 1.25 | NA | NA | NA |
| LAC_PROT20 | 1.27 | NA | 1.31 | NA | 1.38 | NA |
| LAC_BACT10 | 1.76 | NA | NA | NA | NA | NA |
| LAC_BACT11 | 1.37 | NA | 1.32 | 1.32 | NA | NA |
| LAC_IGN07 | 1.29 | NA | 1.30 | 1.48 | 1.39 | 1.41 |
| LAC_VER05 | NA | NA | 1.33 | NA | NA | NA |
| LAC_PROT21 | NA | NA | NA | 1.29 | 1.27 | 1.16 |
| LAC_PROT22 | NA | NA | NA | 1.38 | 1.24 | 1.40 |
| LAC_BAC20 | 1.13 | 1.35 | 1.13 | 1.19 | 1.14 | 1.21 |
| LAC_BACT12 | 1.16 | NA | 1.14 | 1.18 | 1.19 | NA |
| LAC_NIT01 | NA | NA | 1.25 | 1.31 | 1.41 | NA |
| LAC_CHLX06 | 1.24 | NA | 1.21 | 1.28 | 1.18 | 1.28 |
| LAC_PROT22 | 1.41 | NA | 1.36 | 1.24 | 1.23 | NA |
| LAC_MIC04 | NA | NA | NA | 1.35 | 1.23 | NA |
| LAC_PROT23 | NA | NA | NA | 1.47 | NA | NA |
| LAC_PLT02 | NA | 1.56 | 2.13 | 1.07 | 1.13 | 1.12 |
| LAC_CHLX07 | NA | NA | 1.69 | NA | 1.35 | 1.71 |
| LAC_BAC21 | 1.12 | 1.21 | 1.13 | 1.25 | 1.23 | NA |
| LAC_BACT13 | NA | NA | NA | NA | 1.31 | NA |
| LAC_PROT25 | NA | NA | NA | 1.52 | 1.43 | NA |
| LAC_ACD05 | 1.55 | NA | NA | NA | 1.48 | 1.50 |
| LAC_CHLX08 | NA | NA | NA | NA | 1.38 | NA |
| LAC_ARM01 | NA | NA | NA | NA | 1.28 | 1.31 |
| LAC_MIC05 | NA | NA | NA | NA | 1.22 | NA |

| | | | | | | |
|---|---|---|---|---|---|---|
| LAC_PROT26 | NA | NA | NA | NA | 1.29 | 1.26 |
| LAC_CHLX09 | 1.74 | 1.45 | 1.56 | 1.63 | 1.42 | 1.46 |
| LAC_CHLX10 | 1.72 | 1.37 | 1.26 | 1.42 | 1.36 | 1.59 |
| LAC_ACD06 | NA | NA | NA | NA | NA | 1.27 |
| LAC_ACT02 | NA | NA | NA | NA | 1.30 | 1.29 |
| LAC_ACT03 | NA | NA | NA | NA | 1.29 | 1.36 |
| LAC_ACT04 | NA | NA | NA | NA | 1.39 | 1.26 |
| LAC_BAC22 | NA | 1.34 | NA | 1.39 | 1.20 | 1.26 |
| LAC_BAC23 | 1.27 | 1.14 | 1.17 | 1.17 | 1.18 | 1.19 |
| LAC_BAC24 | NA | NA | NA | NA | 1.37 | 1.39 |
| LAC_PROT27 | 2.05 | 2.11 | 2.06 | 1.56 | 1.34 | 1.26 |
| LAC_CHLX11 | NA | NA | NA | NA | NA | 1.29 |
| LAC_CHLX12 | NA | NA | NA | NA | NA | 1.23 |
| LAC_CHLX13 | NA | NA | NA | NA | NA | 1.48 |
| LAC_CHLX14 | NA | NA | NA | NA | NA | 1.33 |
| LAC_CHLX15 | NA | NA | NA | NA | NA | 1.31 |
| LAC_CHLX16 | NA | NA | NA | NA | 1.30 | 1.29 |
| LAC_CHLX17 | NA | NA | NA | NA | 1.35 | 1.25 |
| LAC_PROT28 | NA | NA | 1.33 | 1.27 | 1.20 | 1.19 |
| LAC_CHLX18 | NA | NA | NA | NA | 1.41 | 1.36 |
| LAC_PROT30 | NA | NA | NA | 1.89 | 1.67 | 1.43 |
| LAC_MIC06 | NA | NA | NA | NA | NA | 1.15 |
| LAC_D-T02 | NA | 1.59 | NA | 1.59 | 1.43 | 1.42 |
